# Keratins and Plakin family cytolinker proteins control the length of epithelial microridge protrusions

**DOI:** 10.1101/2020.02.18.954933

**Authors:** Yasuko Inaba, Vasudha Chauhan, Aaron Paul van Loon, Lamia Saiyara Choudhury, Alvaro Sagasti

## Abstract

Actin filaments and microtubules create diverse cellular protrusions, but intermediate filaments, the strongest and most stable class of cytoskeletal elements, are not known to directly participate in the formation of protrusions. Here we show that Keratin intermediate filaments directly regulate the morphogenesis of microridges, elongated protrusions from mucosal epithelial cells arranged in elaborate fingerprint-like patterns. Developing microridges on zebrafish skin cells contained both Actin and Keratin filaments. Keratin filaments maintained microridges upon F-actin disruption, and overexpressing Keratins lengthened microridges. Envoplakin and Periplakin, Plakin family cytolinkers that bind to F-actin and Keratins, localized to microridges and were required for their morphogenesis. Strikingly, Plakin protein levels directly determined microridge length. An actin-binding domain of Periplakin was required to initiate microridge morphogenesis, whereas Periplakin-Keratin binding was required to stabilize and elongate microridges. Our results thus separate microridge morphogenesis into two steps with differential requirements for cytoskeletal elements, expand our understanding of intermediate filament functions, and identify microridges as cellular protrusions that integrate actin and intermediate filaments.

## MAIN TEXT

The three major classes of cytoskeletal elements—microtubules, actin filaments, and intermediate filaments—each have distinct mechanical and biochemical properties, suiting them to different functions. Actin filaments and microtubules create diverse cellular protrusions, such as filopodia, microvilli, and cilia, but intermediate filaments (IFs), despite being the strongest and most stable cytoskeletal elements, are not commonly believed to directly participate in the formation of cellular protrusions.

Microridges are laterally elongated protrusions arranged in striking maze-like patterns on the apical surface of epithelial cells (Depasquale, 2018). They form on a variety of mucosal epithelia in many animals, including the periderm layer of the zebrafish skin, where they are required for maintaining glycans on the skin surface (Pinto et al., 2019). Microridges form from the coalescence of finger-like, actin-based precursor protrusions that we call pegs (van Loon et al., 2020), a process dependent on the F-actin nucleator Arp2/3 (Lam et al., 2015; Pinto et al., 2019; van Loon et al., 2020) and the relaxation of surface tension by cortical myosin-based contraction (van Loon et al., 2020). Although microridges are primarily known as actin-based protrusions, ultrastructural studies have reported the occasional presence of Keratin IFs within microridges (Uehara et al., 1991; Pinto et al., 2019). Here we show that Keratin filaments, in cooperation with actin filaments and Plakin family cytolinkers, play a direct role in microridge morphogenesis.

### Keratins are core components of mature microridges

To investigate Keratin localization in the zebrafish periderm, we tagged six type I Keratin proteins expressed in periderm cells (Cokus et al., 2019) with GFP or mRuby at their C-termini, in bacterial artificial chromosomes (BACs). Imaging periderm cells in live zebrafish expressing these reporters revealed that all Keratins localized in two distinct patterns within a cell: As expected, they formed a filamentous network filling cells; remarkably, they also formed what appeared to be thick bundles in the pattern of microridges at the apical surface (Fig. 1A, S1). Tagging an allele of one of these Keratins, Keratin 17 (Krt17), with CRISPR-facilitated homologous recombination, confirmed that the endogenously expressed protein localized in these two patterns (Fig. 1B).

**Figure 1.**
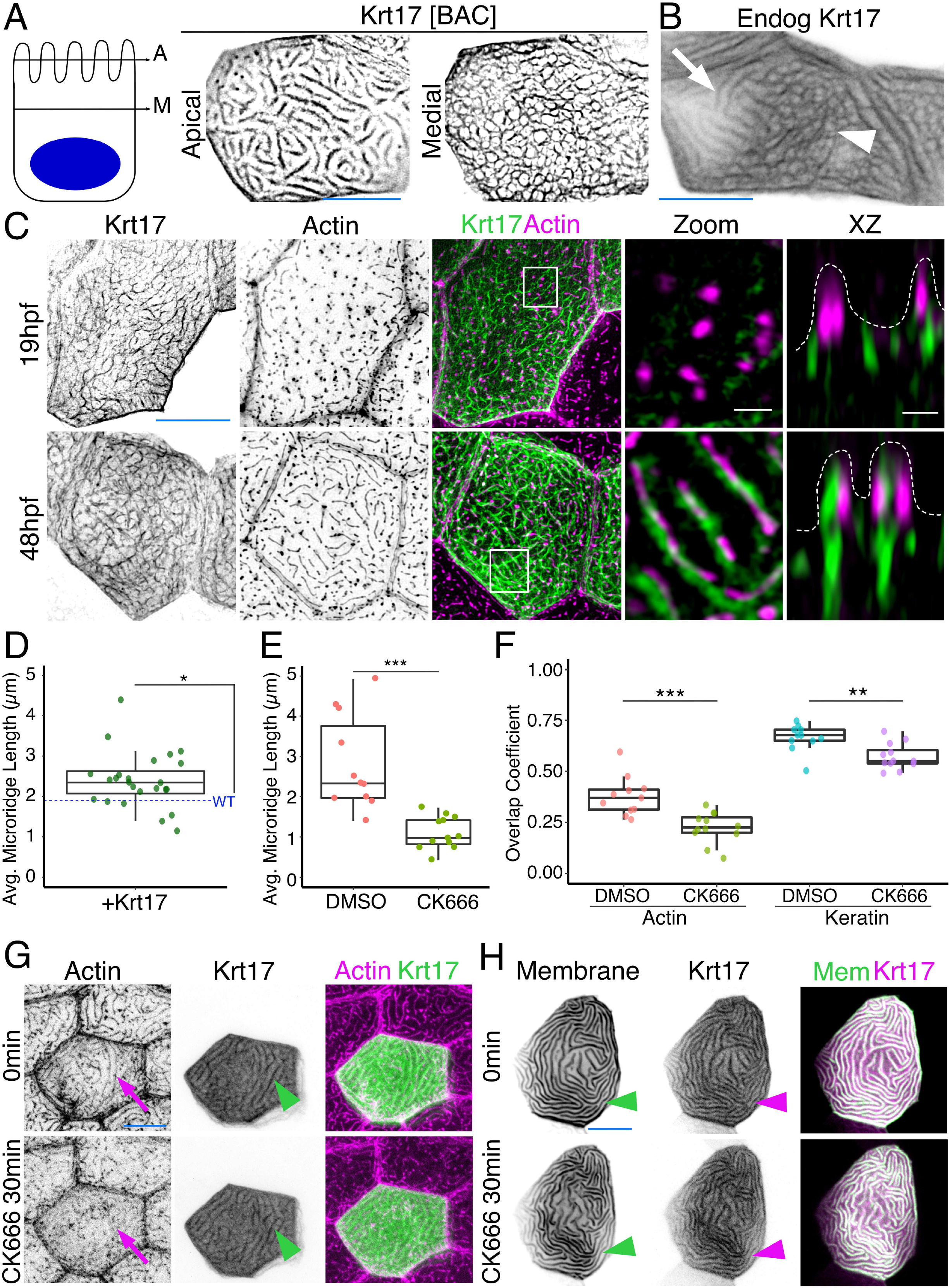
Keratins are core components of microridges. (A) SIM optical sections of a Krt17-GFP[BAC]-expressing zebrafish periderm cell at 48 hpf. Cartoon shows the relative location of Apical (A) and Medial (M) optical sections. (B) Oblique optical section through a cell with endogenously-tagged Krt17. This section shows both the microridge-like pattern at the apical surface of the cell (arrow), and the filamentous pattern deeper in the cell (arrowhead). (C) Projections and orthogonal views of SIM images of Krt17-GFP[BAC]- and Lifeact-Ruby-expressing cells at the indicated developmental stages. White boxes, regions of magnification in zoom panels. Orthogonal views (XZ) at 19hpf show Krt17 is not in pegs, but at 48 hpf localizes alongside Actin in microridges. (D) Dot and box plot of average microridge length per cell in cells over-expressing Krt17-GFP[BAC] at 48hpf. *p<0.05, Wilcoxon rank-sum test. n=24 cells from 4 fish. Blue line shows the median average microridge length per cell in WT animals (see Fig. 3B). (E) Dot and box plot of average microridge length per cell in cells at 48hpf after 30min treatment with DMSO or CK666. ***p<0.005, Wilcoxon rank-sum test. n=11-12 cells from 3 fish. (F) Dot and box plot of microridge and Keratin overlap coefficients, comparing the colocalization of each protein at 0 min with its localization after 30 min DMSO or CK666 treatment. **p<0.01, ***p<0.005, Wilcoxon rank-sum test. n=11-12 cells from 3. (G) Krt17-GFP[BAC]- and Lifeact-Ruby-expressing cells at 48hpf treated with CK666 for 30 minutes. Arrows show that the F-actin microridge pattern was disrupted after 30min of treatment. Arrowheads show that the Krt17 microridge pattern was retained. See Figure S2 for images of control DMSO treatments. (H) GFP-PH-PLC and Krt17-mRuby[BAC] in periderm cells treated with CK666 for 30 minutes at 48hpf. GFP-PH-PLC was expressed by crossing BAC(ΔNp63:Gal4FF)^la213^ to UAS:GFP-PH-PLC, and Krt17-mRuby[BAC] was expressed by transient transgenesis. Arrowheads show that membrane protrusions and the Krt17 microridge pattern were maintained. See Fig. S2 for images of control DMSO treatments, and orthogonal views. Scale bars:10µm (A-C, G-H, blue line) and 1µm (C, white line).

To observe Keratin localization at higher resolution, we used Super-Resolution Structured Illumination Microscopy (SIM) to image cells expressing the Krt17-GFP BAC reporter. At an early stage of microridge morphogenesis (19 hours post-fertilization, hpf), when periderm surfaces are dominated by pegs (van Loon et al., 2020), Krt17 formed the filamentous pattern in cells, but did not localize within pegs. At a later stage (48 hpf), when mature microridges have formed, Krt17 invaded microridges and appeared to form filaments alongside F-actin (Fig. 1C). Intriguingly, average microridge length per cell, as measured by imaging the Actin reporter Lifeact-Ruby, was longer in Krt17-GFP over-expressing cells than in wild-type (WT) cells (Fig. 1D). These observations confirm that Keratins are components of mature microridges, and suggest that they may play a role at later stages of microridge morphogenesis.

### Keratins are stable microridge components

Since IFs are the strongest and most stable of the cytoskeletal filaments, we wondered if they might contribute to microridge stability. To destabilize F-actin in microridges, we treated animals for 30 minutes with the Arp2/3 inhibitor CK666, which causes microridges to disassemble and revert back into peg-like structures (Fig. 1E) (Lam et al., 2015; Pinto et al., 2019; van Loon et al., 2020). Strikingly, despite the loss of the F-actin microridge pattern, the apical Krt17-GFP microridge pattern was retained (Fig. 1F-G, S2A). Labeling cells with a membrane reporter (GFP-PH-PLC) revealed that overexpressing the Krt17-GFP BAC reporter preserved the protrusive membrane topology upon F-actin disruption (Fig. 1H, S2B-C). These results support the idea that Keratins play roles in stabilizing and/or elongating microridges at a later stage of morphogenesis.

### The cytolinker proteins Envoplakin and Periplakin localize to microridges

If F-actin and Keratins both contribute to microridge morphogenesis, we speculated that they may interact via linker proteins. By examining larval periderm cell transcriptomes (Cokus et al., 2019), we identified two cytolinker proteins, Envoplakin (Evpl) and Periplakin (Ppl), which are highly expressed and enriched in periderm cells, relative to other epithelial cells. Evpl and Ppl, members of the Plakin protein family (Jefferson et al., 2004), contribute to keratinization of the mammalian skin (Ruhrberg et al., 1997, 1996), heterodimerize with each other (Kalinin et al., 2004), and can bind both F-actin (Kalinin et al., 2005) and Keratins (Kazerounian et al., 2002; Karashima and Watt, 2002), thus potentially linking the two types of cytoskeletal filaments in microridges. To determine the localization of Evpl and Ppl in zebrafish periderm cells, we made Ppl-GFP, Evpl-GFP, and Evpl-mRuby BAC reporter fusions and imaged them in transient transgenic animals. Evpl and Ppl both localized to microridges (Fig. 2). Similar to its behavior in mammalian cell culture (DiColandrea et al., 2000), when expressed on its own, Evpl formed prominent aggregates, which were reduced when Ppl was co-expressed (Fig. 2A-B, S3), consistent with the possibility that the two Plakin proteins dimerize or oligomerize. We verified the Ppl localization pattern by GFP-tagging the endogenous gene with CRISPR-facilitated homologous recombination (Fig. 2C). SIM microscopy of the co-expressed BAC reporters revealed that Evpl and Ppl localize within microridges. In optical sections along the z-axis, Ppl and Evpl appeared to be adjacent to F-actin and Keratin filaments, but were almost completely overlapping with each other (Fig. 2D). In an x-y section, however, Evpl and Ppl were arranged in an apparently alternating pattern, consistent with their ability to assemble into a higher order oligomeric arrangement (Kalinin et al., 2004) (Fig. 2E).

**Figure 2.**
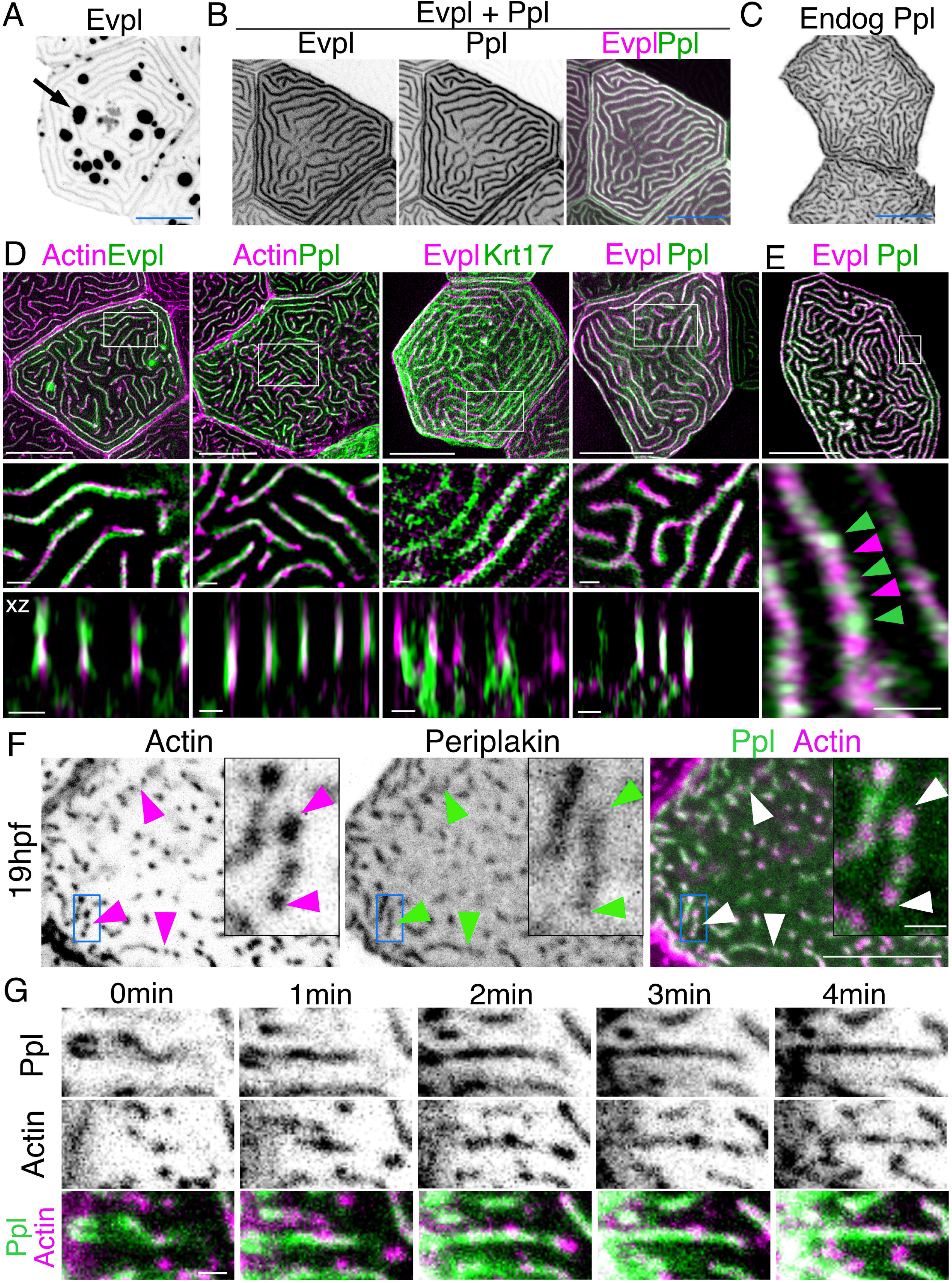
Evpl and Ppl localize to microridges and are required for initiation of microridge morphogenesis. (A) Evpl-mRuby[BAC] expression in periderm cells of 48hpf zebrafish showing localization in aggregates (e.g. arrow) and microridges. (B) Evpl-mRuby[BAC] and Ppl-GFP[BAC] co-expression in a periderm cell at 48hpf. (C) Endogenously tagged Ppl-GFP expression in periderm cells at 48hpf. (D) Projection and orthogonal views of SIM images of the indicated co-expressed reporters. White boxes, regions of magnification in middle panels. Bottom panel, orthogonal view (apical up, basal down). (E) Confocal airyscan image of Ppl-GFP[BAC] and Evpl-mRuby[BAC]. White box, region of magnification in lower panel. Arrowheads show alternating arrangement of Ppl and Evpl in microridges. (F) Ppl-GFP[BAC] and Lifeact-Ruby expression in a 19hpf periderm cell. Arrowheads point to representative areas in which Ppl localizes to longer structures than actin pegs. Boxes, area of magnification for insets. (G) Sequential projections from a time-lapse movie of Ppl-GFP[BAC] and Lifeact-Ruby at the end of cytokinesis (24hpf). Note that Ppl structures appear to precede F-actin in forming microridges. Scale bars: 10µm (A-F) and 1µm (zoomed and orthogonal views in D, F, and G).

To determine when Plakins first localize to microridges, we observed Ppl localization at an earlier stage (19 hpf), when periderm cell surfaces are dominated by pegs. Unlike Keratin, at this stage, Ppl associated with pegs and, surprisingly, often appeared to form longer and more continuous structures than F-actin itself (Fig. 2F). To observe their localization with greater temporal precision, we imaged Ppl and F-actin reporters in time-lapse movies of cells undergoing cytokinesis. Just before periderm cell division, microridges dissolve, but then re-assemble at the end of cytokinesis (Lam et al., 2015), thus affording us an opportunity to image the entire process with rapid, predictable kinetics. These movies revealed that Ppl forms elongated, continuous structures immediately before the coalescence of F-actin pegs into microridges, potentially as part of a template for microridge assembly (Fig. 2G). Since Keratins do not localize to microridges until later in development, these observations suggest that Ppl likely plays a Keratin-independent role in the initiation of microridge morphogenesis.

### Envoplakin and Periplakin determine microridge length

To determine the function of Evpl and Ppl in microridge morphogenesis, we created stable zebrafish mutant lines by deleting several exons of each gene with the CRISPR/Cas9 system (Fig. S4). At larval stages, *ppl* mutants lacked microridges, forming only pegs, whereas *evpl* mutants formed pegs and short microridges (Fig. 3A-B). Periderm cells in double mutants were indistinguishable from those in *ppl* mutants, projecting only pegs. Defects in microridge organization persisted into adulthood in both mutants (Fig. S5), but all single and double mutant animals were homozygous viable and fertile. Unexpectedly, in both *evpl* and *ppl* heterozygotes, average microridge length per cell was shorter than in WT cells, but longer than in homozygous mutants (Fig. 3B). Average microridge length was shorter in trans-heterozygous mutants (*evpl^+/−^*; *ppl^+/−^*) than in either heterozygous mutant alone, but longer than in homozygous mutants. These observations reveal that *evpl* and *ppl* mutants are semi-dominant and that the dose of Plakin proteins dictates microridge length.

**Figure 3.**
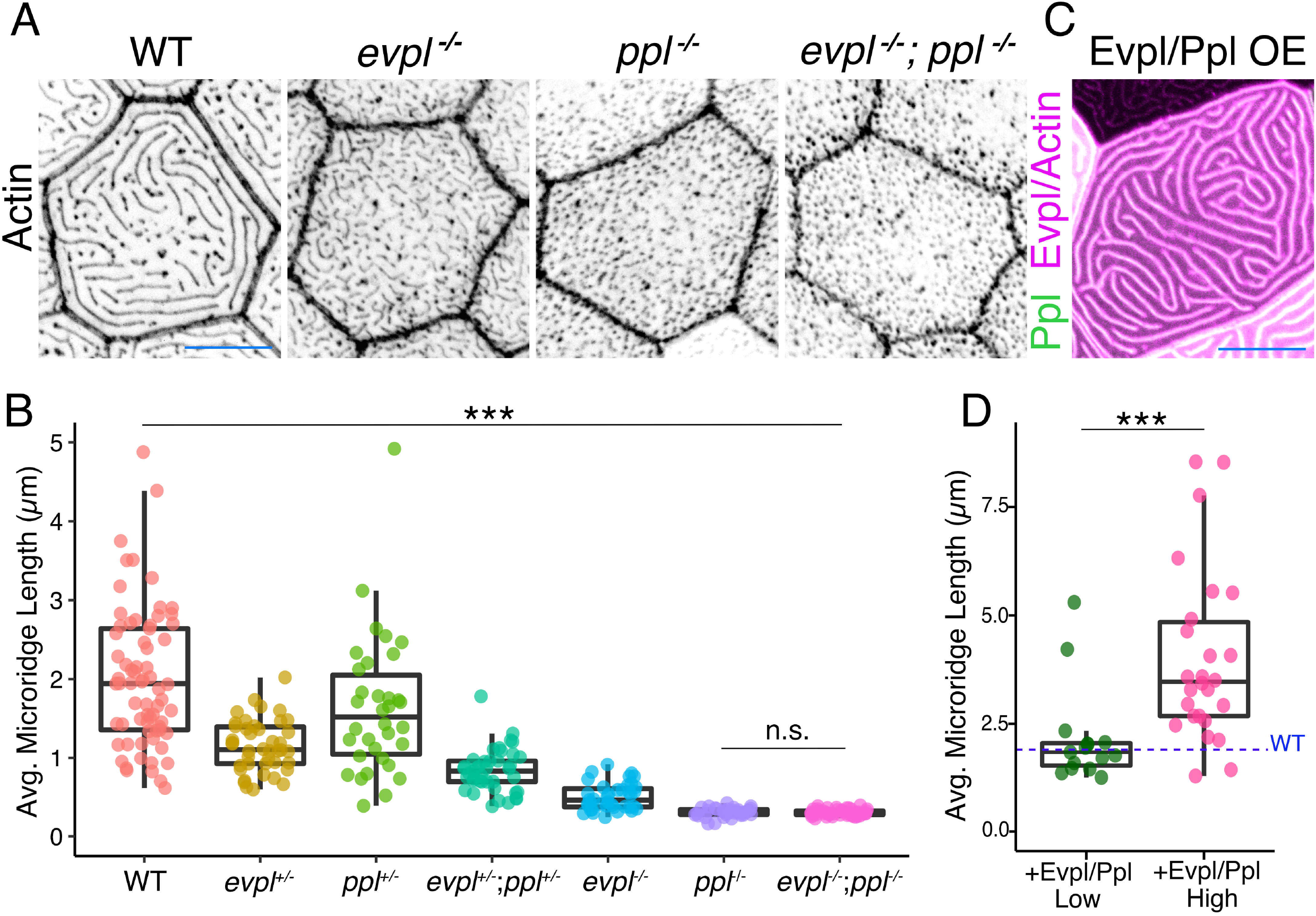
Evpl and Ppl are required for microridge morphogenesis and determine their length. (A) Actin (Lifeact-Ruby) in WT, *evpl*^−/−^, *ppl*^−/−^ and *evpl^−/−^;ppl^−/−^* mutants at 48hpf. (B) Dot and box plot of average microridge length per cell at 48hpf from animals of the indicated genotypes. All comparisons between each genotype were significantly different from one another, except where indicated as n.s. ***p<0.0001, Wilcoxon rank-sum test. n=27-69 cells from 3-9 fish per genotype. (C) Cell overexpressing Evpl-mRuby[BAC], Ppl-GFP[BAC] and Lifeact-Ruby at 48hpf. (D) Dot and box plot of average microridge length per cell in cells over-expressing Evpl-mRuby[BAC] and Ppl-GFP[BAC] at 48hpf. See Figure S6 for categorization into “low” and “high” overexpression groups. ***p<0.0001, Wilcoxon rank-sum test. n=41 cells from 5 fish were analyzed. Blue line shows the median average microridge length per cell in WT animals (from B). Scale bars: 10µm(A and C).

To determine if Evpl and Ppl are not only required, but also sufficient for lengthening microridges, we co-overexpressed the Evpl-mRuby and Ppl-GFP BAC reporters in WT animals (Fig. 3C). Since the reporters were fluorescently tagged, we could estimate the relative concentration of overexpressed Plakins in each cell from its fluorescence intensity. Plotting fluorescence intensity versus average microridge length per cell revealed that cells expressing higher levels of the Plakins tended to have longer microridges (Fig. S6). Indeed, grouping cells into high- and low-expressing categories demonstrated that microridges in cells with high Plakin levels were substantially longer than microridges in WT cells (Fig. 3D, S6). Together, these results indicate that Ppl and Evpl function like a molecular rheostat for microridge length: lowering Plakin expression shortens microridges, whereas increasing Plakin expression lengthens microridges.

### Plakins associate with microridges and Keratins in zebrafish skin cells

Given their ability to bind F-actin and Keratins (Kazerounian et al., 2002; Karashima and Watt, 2002; Kalinin et al., 2005), we hypothesized that Evpl and Ppl serve as linkers between Keratins and F-actin in microridges. Previous biochemical studies showed that their N-terminal head domains can bind to Actin (and potentially also membranes) (Kalinin et al., 2005), their rod domains promote dimerization (Kalinin et al., 2004; DiColandrea et al., 2000) and their C-terminal tail domains bind to Keratins (Kazerounian et al., 2002; Karashima and Watt, 2002). To determine the localization of these domains in zebrafish periderm cells, we GFP- or tdTomato-tagged each domain of each Plakin protein and expressed them in periderm cells of WT animals. As expected, full-length Plakins localized to microridges, but the isolated head and tail domains of each Plakin localized throughout the cytoplasm and nucleus, suggesting that dimerization via the rod domain is required for their localization to microridges (Fig. 4A). The Ppl rod domain, which is required for dimerization in vitro (Kalinin et al., 2004), localized variably to microridges. However, the Evpl rod domain did not localize to microridges (Fig. 4A), suggesting that it is not sufficient for dimerization, and implying that the Ppl rod domain may weakly homodimerize, as previously suggested by biochemical studies (Kalinin et al., 2004). Indeed, whereas full-length Ppl localized to the short microridges in Evpl mutants, full-length Evpl was cytoplasmic in Ppl mutants (Fig. 5), consistent with the hypothesis that Ppl, but not Evpl, can homodimerize (or homo-oligomerize), and that dimerization is required for microridge localization. This observation may also explain why Ppl mutants have a more severe microridge phenotype than Evpl mutants (Fig. 3).

**Figure 4.**
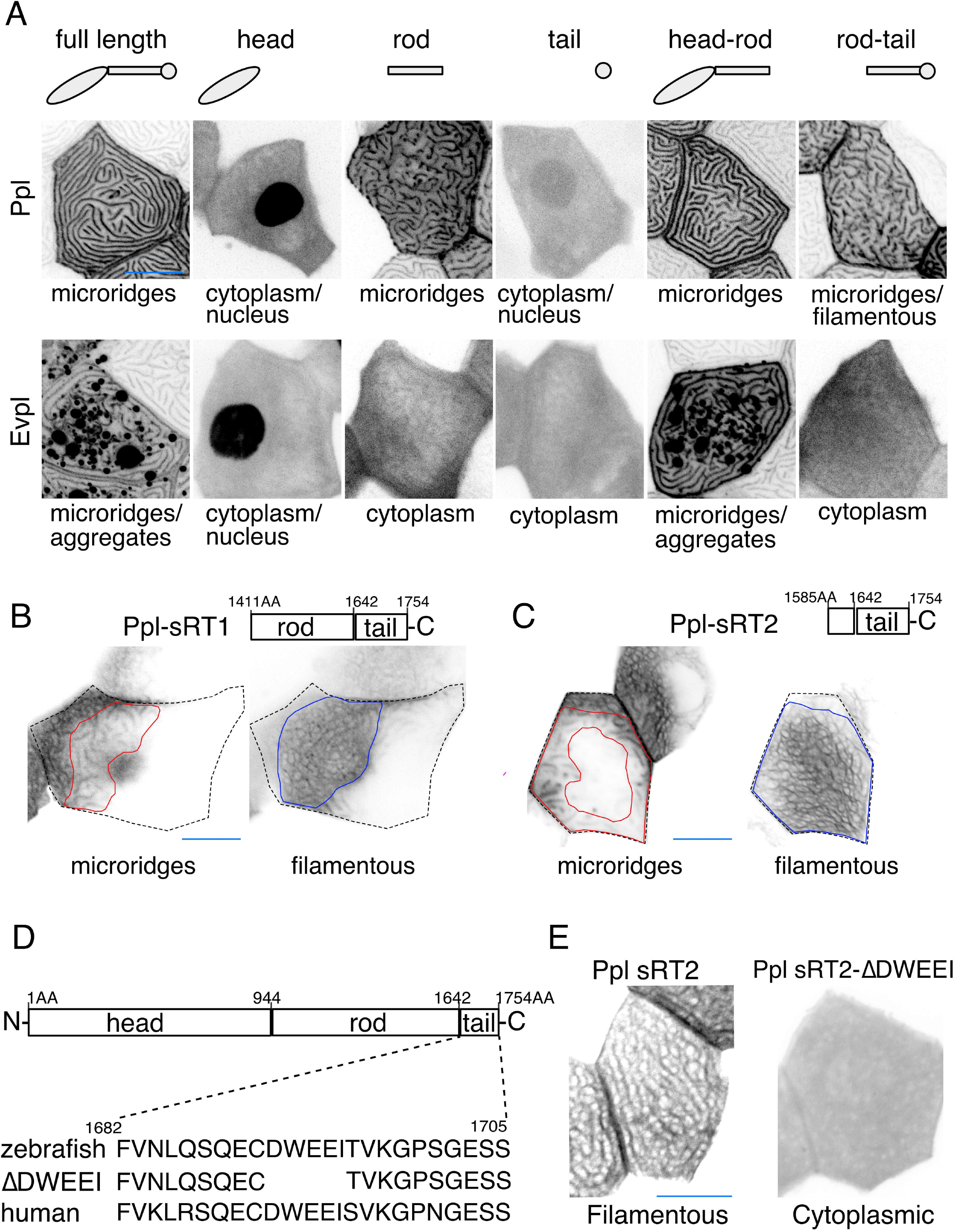
Ppl and Evpl domain localization. (A) Projections of Evpl-tdTomato and Ppl-GFP variants expressed in periderm cells at 48hpf. Schematic indicates the domains in each variant. (B-C) Optical sections of GFP-tagged Ppl truncated rod-tail fusions expressed in periderm cells at 48hpf. Sections show primarily the microridge-like pattern at the apical surface of the cell (left, red outlines), or the filamentous pattern deeper in the cell (right, blue outlines). Top: Diagram of Ppl protein domains. Amino acid numbers are indicated. (D) Top: Diagram of Ppl protein domains. Amino acid numbers are indicated above. Bottom: Amino acid sequence of IF-binding domain of zebrafish and human Ppl, showing the location of the ΔDWEEI mutation. (E) Localization of tagged Ppl(sRT2) and Ppl(sRT2-ΔDWEEI) expressed in WT animals. Scale bars: 10µm (A-C, and E)

**Figure 5.**
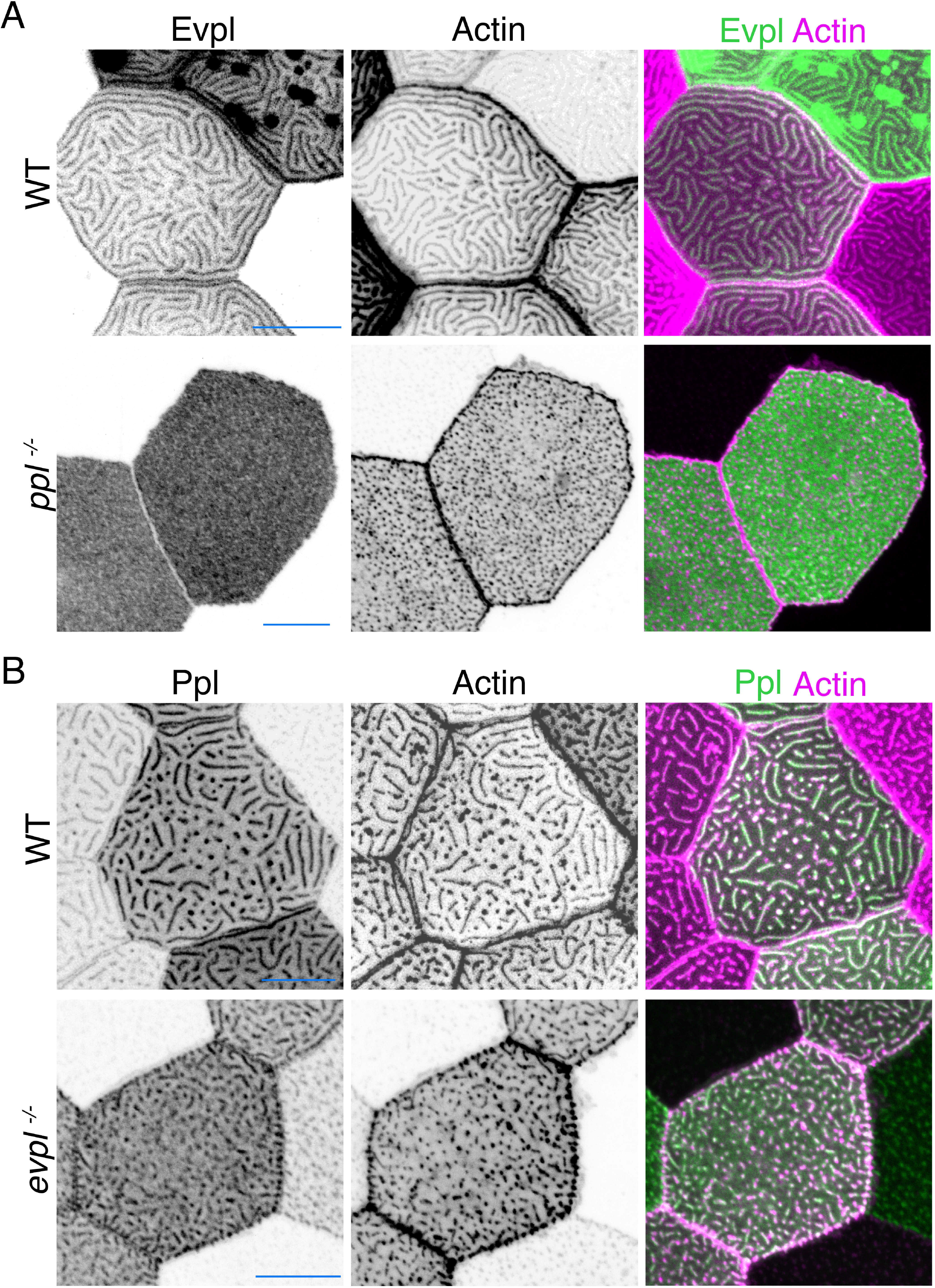
Evpl depends on Ppl for apical localization, but not vice versa. (A) Projection images of Evpl-GFP[BAC] and Actin (Lifeact-Ruby) expressed in WT and *ppl^−/−^* periderm cells at 48hpf. Evpl localized to microridges in WT and to the cytoplasm in *ppl^−/−^* periderm cells. (B) Projection images of Ppl-GFP[BAC] and Actin (Lifeact-Ruby) expressed in WT and *evpl^−/−^* periderm cells. Ppl localized to microridges in WT and into pegs/short microridges in *evpl^−/−^* periderm cells at 48hpf. Scale bars,10µm.

To determine where dimerized head and tail domains localize, we expressed GFP-tagged head-rod (i.e. Δtail) or rod-tail (i.e. Δhead) domain fusions. Head-rod fusions of both Plakins localized to microridges more robustly than rod domains alone, suggesting that the dimerized head domain enhances localization to microridges, potentially through F-actin binding. Ppl, but not Evpl, rod-tail domain fusions variably localized in Keratin filament-like patterns, suggesting that the dimerized Ppl tail domain associates with Keratins (Fig. 4A). Since parts of the rod domain can contribute to F-actin binding (Kalinin et al., 2005), we made two shorter fusions containing only part of the Ppl rod domain and the entire Ppl tail domain (sRT1-GFP and sRT2-GFP). These shorter fusions strongly localized in Keratin-like patterns that included the thick microridge-associated bundles and the filamentous network throughout the cell (Fig.4B-C). Together these findings demonstrate that dimerized Plakins have the potential to link F-actin with Keratins in microridges.

### The Periplakin head domain is required at an early step of morphogenesis to fuse pegs into microridges

To determine how each Plakin domain contributes to microridge morphogenesis, we attempted to rescue Plakin mutants with full-length and truncated tagged versions of the Plakins. Injecting genes encoding fluorescently tagged full-length Evpl-tdTomato and Ppl-GFP into *evpl^−/−^*;*ppl^−/−^* double mutant fish rescued microridge development. Strikingly, average microridge length per cell correlated with fluorescence intensity (Fig. 6A-B), further illustrating that Plakin expression levels determine microridge length. By contrast, truncated fusion proteins lacking the Plakin head domains did not rescue microridge length in double mutants (Fig. 6A-B). Similarly, Ppl rod-tail fusions did not rescue Ppl single mutants (Fig. S7), suggesting that heterodimers containing only the Envoplakin head domain cannot support microridge morphogenesis.

**Figure 6.**
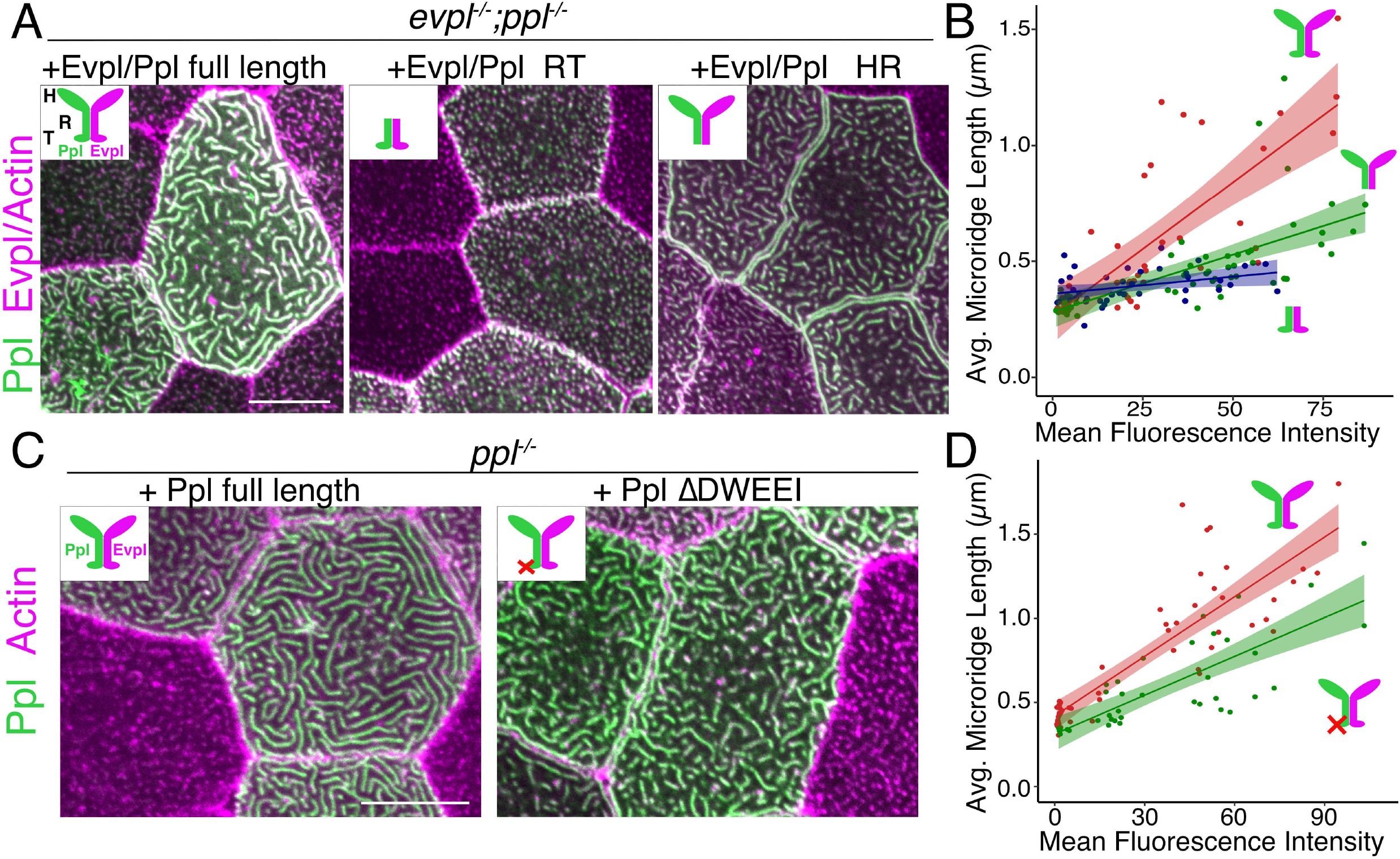
The Periplakin head domain is required for initiation of microridge morphogenesis, and its tail domain is required to elongate microridges. (A, C) Lifeact-Ruby-expressing cells mosaically expressing Ppl-GFP and Evpl-tdTomato variants in periderm cells. Neighboring cells lacking GFP served as controls. (A) Periderm cells of *evpl^−/−^;ppl^−/−^* mutant animals co-expressing full-length tagged Evpl and Ppl (left), tagged Evpl and Ppl rod-tail fusions (middle) or tagged Evpl and Ppl head-rod fusions (right). Inset schematics show the domains expressed. H: Head; R: Rod; T: Tail domain. (B) Scatter plot of average microridge length versus mean fluorescence intensity in *evpl^−/−^;ppl^−/−^* mutant periderm cells expressing full-length Evpl and Ppl (red line), Evpl and Ppl head-rod fusions (green line), or Evpl and Ppl rod-tail fusions (purple line). Shading indicates 95% confidence interval. Slopes=0.0115, 0.0049 and 0.0015; R2= 0.56, 0.44 and 0.13 for red, green and purple lines, respectively. n=39-55 cells from 3-4 fish were analyzed per category. (C) Periderm cells of *ppl^−/−^* mutant animals expressing tagged WT Ppl (left) or the Ppl(ΔDWEEI) mutant (right). (D) Scatter plot of average microridge length versus mean fluorescence intensity in *ppl^−/−^* periderm cells expressing full-length Ppl (red line) or Ppl(ΔDWEEI) (green line). Shading indicates 95% confidence interval. Slopes=0.0118 and 0.0077; R=0.74 and 0.6 for red and green line, respectively. n=37-51 cells from 3-4 fish were analyzed per category. Scale bars: 10µm (A, and C).

### The Keratin-binding Periplakin tail domain is required to elongate microridges

To ask if Plakin-Keratin interactions play a role in microridge morphogenesis, we expressed tagged Evpl and Ppl head-rod domain fusions, which lack the Keratin-binding tail domain, in double mutant embryos. These fusions rescued the initiation of microridge formation, but did not rescue microridge length as well as full-length Plakins (Fig. 6A-B). These results indicate that the Evpl and Ppl dimerized head domains are sufficient for the initiation of microridge morphogenesis, but not for their full elongation. Similarly, expressing the Ppl head-rod fusion in Ppl single mutants allowed the formation of only short, but not fully elongated, microridges (Fig. S7). Since the Plakin tail domains bind to Keratins, these experiments suggest that Plakin-Keratin association may facilitate microridge elongation.

To further test if Plakin-Keratin interactions promote microridge elongation, we deleted five amino acids in the Ppl tail domain required for Keratin binding (ΔDWEEI) (Fig. 4D) (Karashima and Watt, 2002). Localization of Ppl(ΔDWEEI) rod-tail fusions (sRT2(ΔDWEEI)-GFP) to the Keratin network was severely reduced (Fig. 4E), confirming *in vivo* that this site is required for optimal Ppl-Keratin binding. Similar to head-rod fusions, Ppl(ΔDWEEI) could not rescue microridge length as well as WT Ppl (Fig. 6C-D). Deleting another small domain required for Keratin biding (Δ1/2Box2) (Karashima and Watt, 2002) in Ppl yielded similar results (not shown). These results suggest that Ppl-Keratin interactions are required for full microridge elongation.

### Plakin-Keratin interactions stabilize microridges

IFs are the strongest cytoskeletal elements (Janmey et al., 1991), maintain their structure even as they replace subunits (Çolakoğlu and Brown, 2009), and protect cells from mechanical stress (Leube et al., 2017). We therefore hypothesized that Plakin-mediated recruitment of Keratins into microridges stabilizes them, and that stabilization permits their elongation. Notably, the IF Vimentin has analogously been hypothesized to play a role in the growth of invadopodia at a late morphogenetic step (Schoumacher et al., 2010). To test our hypothesis, we first compared the stability of short, nascent microridges in early development (24 hpf), when Keratins are not found abundantly in microridges, to their stability at a later stage (48 hpf), when microridges are longer and contain Keratins. Over the course of 5-minute time-lapse movies, the microridge pattern changed more rapidly at 24 hpf than at 48 hpf, indicating that shorter microridges likely lacking Keratins are less stable than longer Keratin-containing microridges in older animals (Fig. 7A-B). To test if Plakin-Keratin binding plays a role in stabilization, we compared cells in double mutant animals expressing full-length Evpl and Ppl to those expressing the Evpl and Ppl head-rod domains (i.e. Δtail domain) at 48 hpf. Microridges were more dynamic in cells expressing Evpl and Ppl head-rod fusions than those expressing full-length Plakins (Fig. 7C-D), similar to microridges in WT cells of younger animals. These results suggest that Plakin-Keratin interactions stabilize microridges to allow them to elongate.

**Figure 7.**
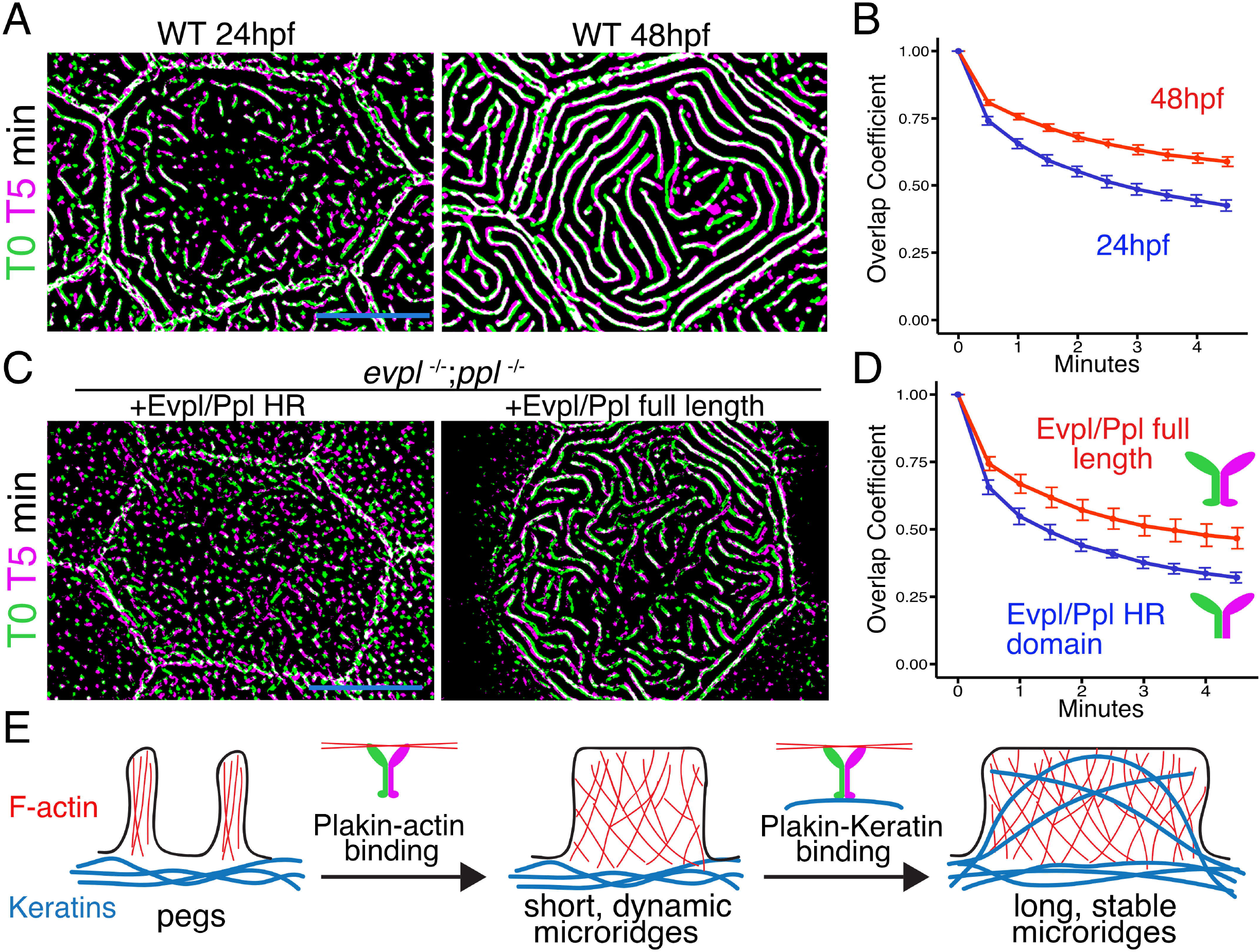
The Periplakin tail domain is required to stabilize microridges. (A) Superimposition of first and last frames from a time-lapse movie of WT cells expressing Lifeact-GFP at the indicated developmental stages. Green: Starting time point, Magenta: 5-minute time point. Overlap is white. (B) Line plots of microridge overlap coefficients at the indicated stages, comparing each time point to the first. n=21-33 cells from 4-5 fish per category. (C) Superimposition of first and last frames from a time-lapse movie, as in E, of *evpl^−/−^;ppl^−/−^* double mutant animals rescued with the indicated Evpl and Ppl fusions. (D) Line plots of microridge overlap coefficients in cells expressing the indicated Evpl and Ppl fusions, comparing each time point to the first. n=11 cells from 3-4 fish were analyzed per category. (E) Two-step model of microridge morphogenesis. First, Plakins interact with F-actin via their head domains to fuse pegs into short microridges. Second, Plakin-Keratin interactions through the Ppl tail domain stabilize microridges, allowing them to further elongate. Scale bars: 10µm (A and C).

In summary, our results reveal that Plakins contribute to microridge morphogenesis at two distinct steps (Fig. 7E). At an early stage, Evpl and Ppl promote the coalescence of pegs into nascent microridges, a process that also requires Arp2/3 and cortical contraction (Lam et al., 2015; Pinto et al., 2019; van Loon et al., 2020). Keratin is not present in microridges at this initial stage, and this step requires the Plakin N-terminal domains, which can bind F-actin (Kalinin et al., 2005), but not their Keratin-binding C-terminal domains. These observations indicate that this early morphogenetic step involves a Keratin-independent Plakin function that could include F-actin interactions. By contrast, Keratin is present in longer, more stable microridges, and the Keratin-binding sites of Plakins are required for full microridge elongation. Since IFs are known for their strength and stability, and Plakin-Keratin binding is required for microridge stabilization and elongation, we propose that the Plakins recruit Keratin into microridges to permit their elongation.

A landmark study comparing the three classes of cytoskeletal elements speculated that the mechanical “differences between F-actin and vimentin [a Keratin-related intermediate filament] are optimal for the formation of a composite material with a range of properties that cannot be achieved by a single polymer network” (Janmey et al., 1991). By combining these two cytoskeletal elements, microridges may achieve an optimal balance between plasticity and stability.

## MATERIALS AND METHODS

### Zebrafish

Zebrafish (Danio rerio) were grown at 28.5°C on a 14 h/10 h light/dark cycle. Embryos were raised at 28.5°C in embryo water (0.3g/L lnstant Ocean Salt, 0.1 % methylene blue). For live confocal imaging, pigmentation was blocked by treating embryos with phenylthiourea (PTU) at 24hpf. All experimental procedures were approved by the Chancellor’s Animal Research Care Committee at UCLA.

### CRISPR/Cas9 mutagenesis

To generate guide RNAs (gRNAs) we used the “Short oligo method to generate gRNA”, as previously described (Talbot and Amacher, 2014). Two Cas9 binding sites were selected for each gene. *Evpl* targeting sequences were located in exons 3 and 7, *ppl* targeting sequences were in exons 2 and 5. DNA template was PCR amplified to make a product containing a T7 RNA polymerase promoter, the gene targeting sequence, and a gRNA scaffold sequence. PCR products were used as a template for RNA synthesis with T7 RNA polymerase (New England Biolabs) and purified (QIAGEN RNA purification kit) to generate gRNAs. Injection mixes contained Cas9 protein (1 mg/ml) (IDT), gRNAs (0.5-1 ng/ul) and 300mM KCI. Injection mixes were incubated on ice 15 min before injection. Embryos were injected at the single-cell stage with 2-5 nl of injection mix. To identify germline founders, F0 founder fish were crossed to wild-type fish and 48hpf embryos were collected for PCR genotyping. Founder progeny were raised to adulthood to establish stable mutant lines.

### BAC reporters

Tg(Krt5:Lifeact-GFP) and Tg(Krt5:Lifeact-mRuby) lines were previously described (van Loon et al., 2020). To create translational fusion transgenes, GFP or mRuby reporter gene cassettes were recombined into a site directly preceding the stop codon of target genes in Bacterial Artificial Chromosomes (BACs), as previously described (Suster et al., 2011; Cokus et al., 2019). BAC identifiers are listed in Table 3.

**Table 1.**
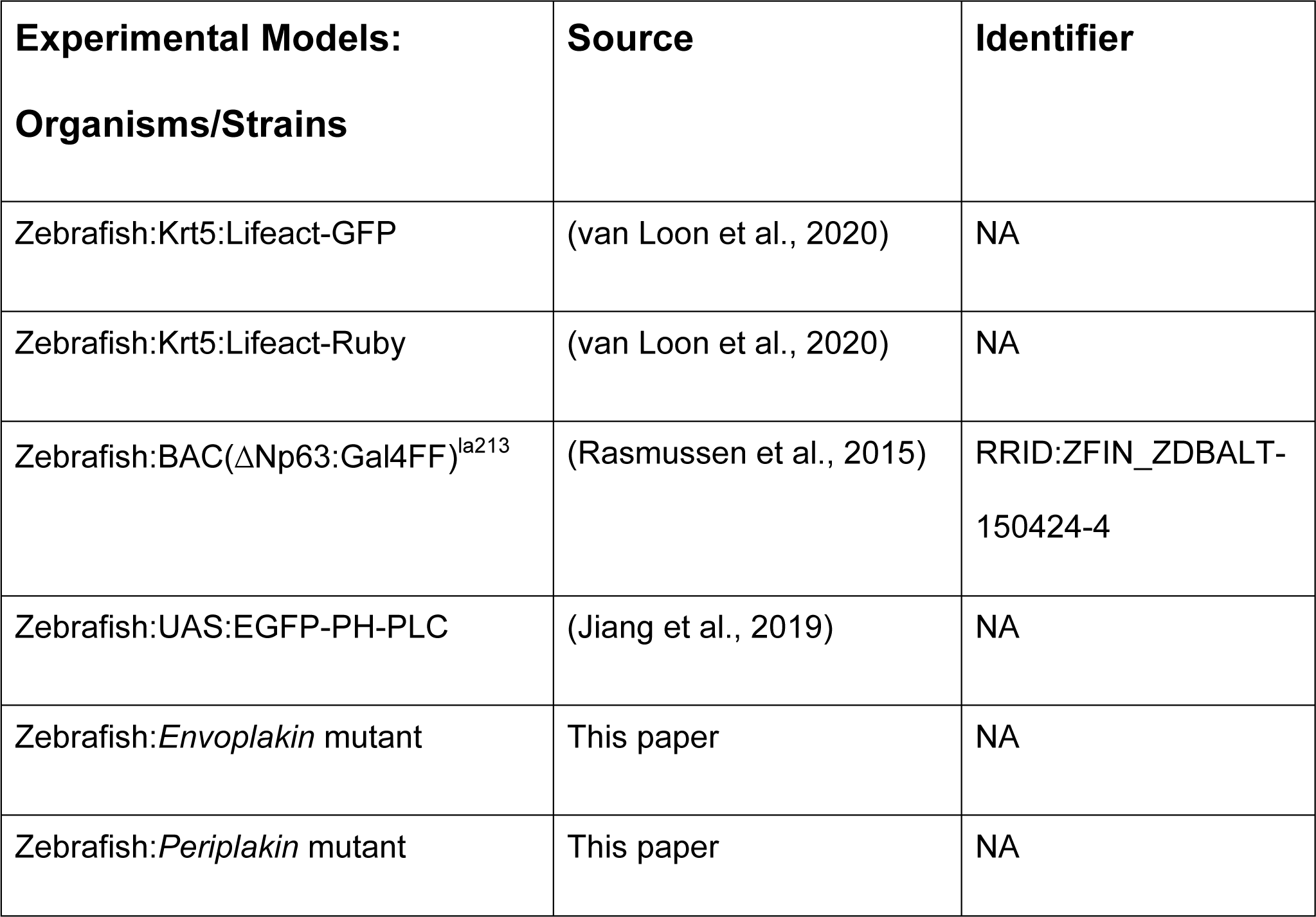
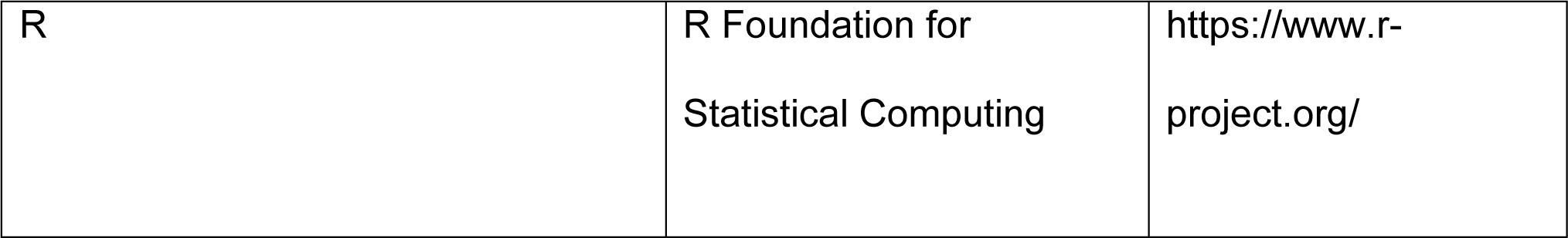
Reagents and resources.

**Table 2.**
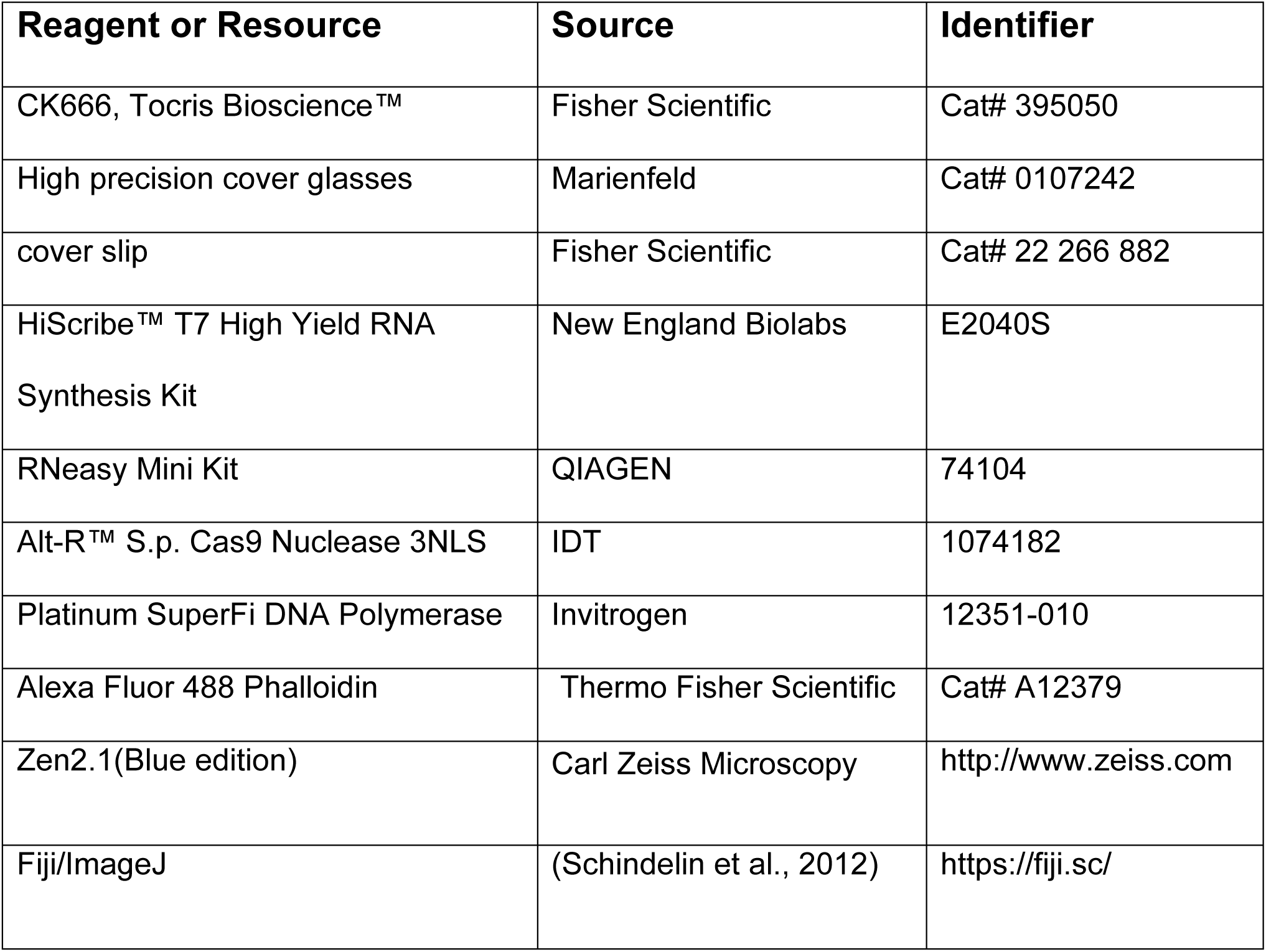
Zebrafish lines.

**Table 3.** Bacterial Artificial Chromosomes.

| **Gene** | **Ensembl#** | **BAC#** |
| --- | --- | --- |
| Envoplakin | ENSDARG00000019808 | Ch211-188J19 |
| Periplakin | ENSDARG00000101043 | Ch73-360K2 |
| Krt17 | ENSDARG00000094041 | Ch73-107M18 |
| Krt99 | ENSDARG00000019365 | Ch73-294H4 |
| Krt92 | ENSDARG00000036834 | Ch73-107M18 |
| krt1-19d | ENSDARG00000023082 | Ch73-107M18 |
| cyt1 | ENSDARG00000092947 | Ch73-107M18 |
| cyt1l | ENSDARG00000036832 | Ch73-107M18 |

### Plasmids

Plasmids were constructed using the Gateway-based Tol2kit (Kwan et al., 2007). Primer sequences are listed in Table 4. The following plasmids were previously described: p5E-Krt5(Rasmussen et al., 2015), pME-EGFPpA, p3E-EGFPpA, p3E-tdTomato and pDestTol2pA2 (Kwan et al., 2007), Krt5-Lifeact-GFP and Krt5-Lifeact-Ruby (van Loon et al., 2020).

**Table 4.**
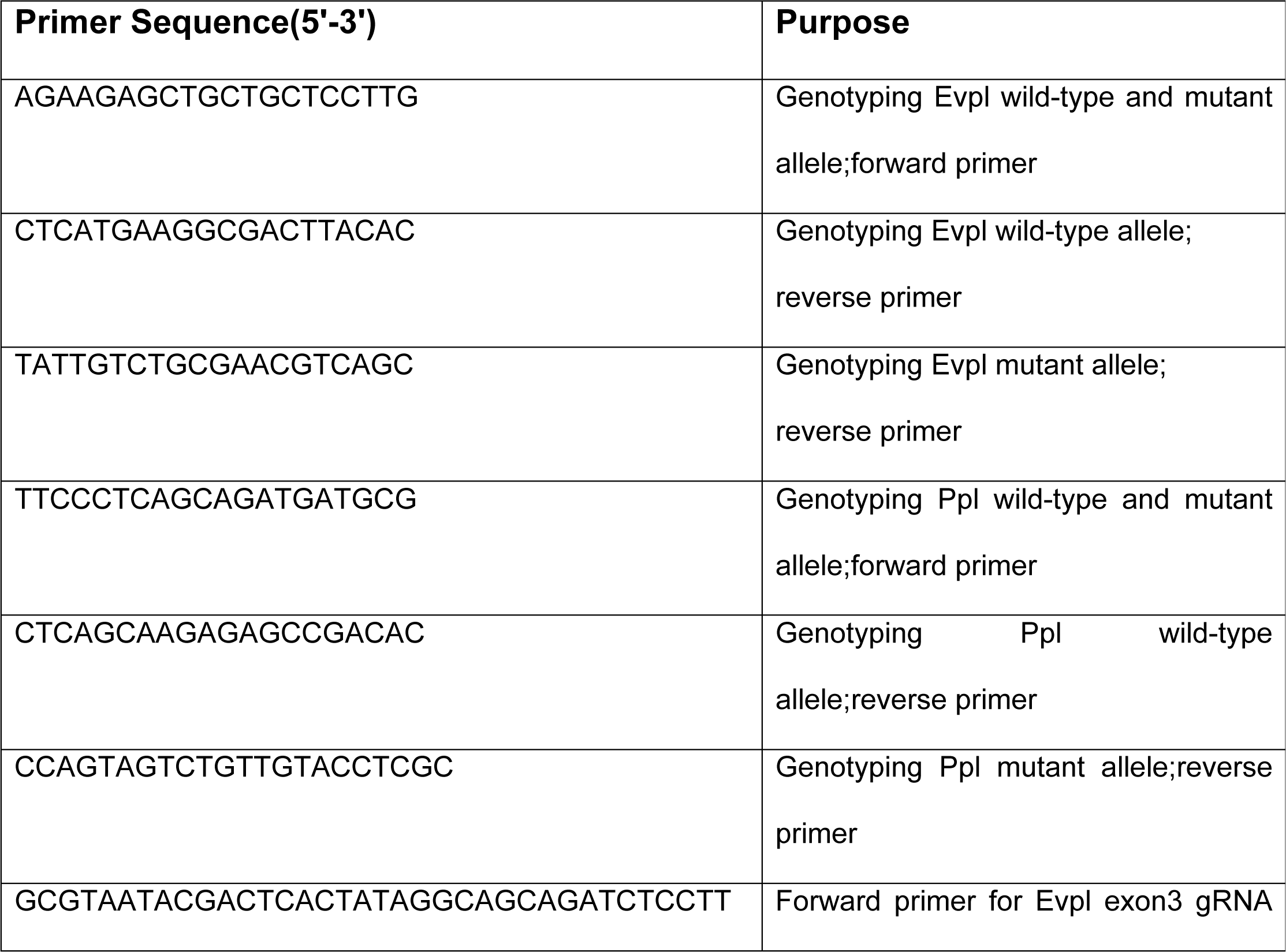
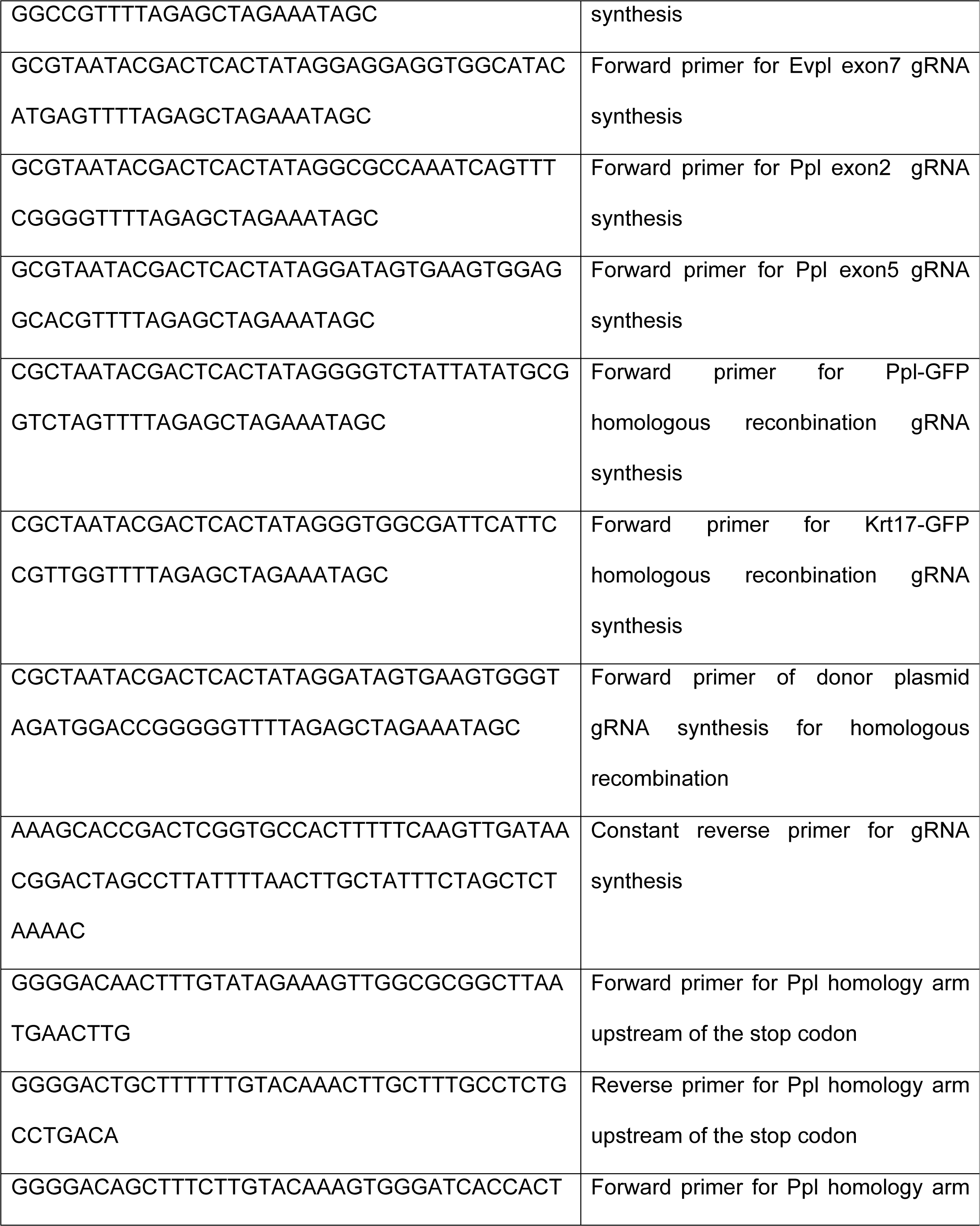
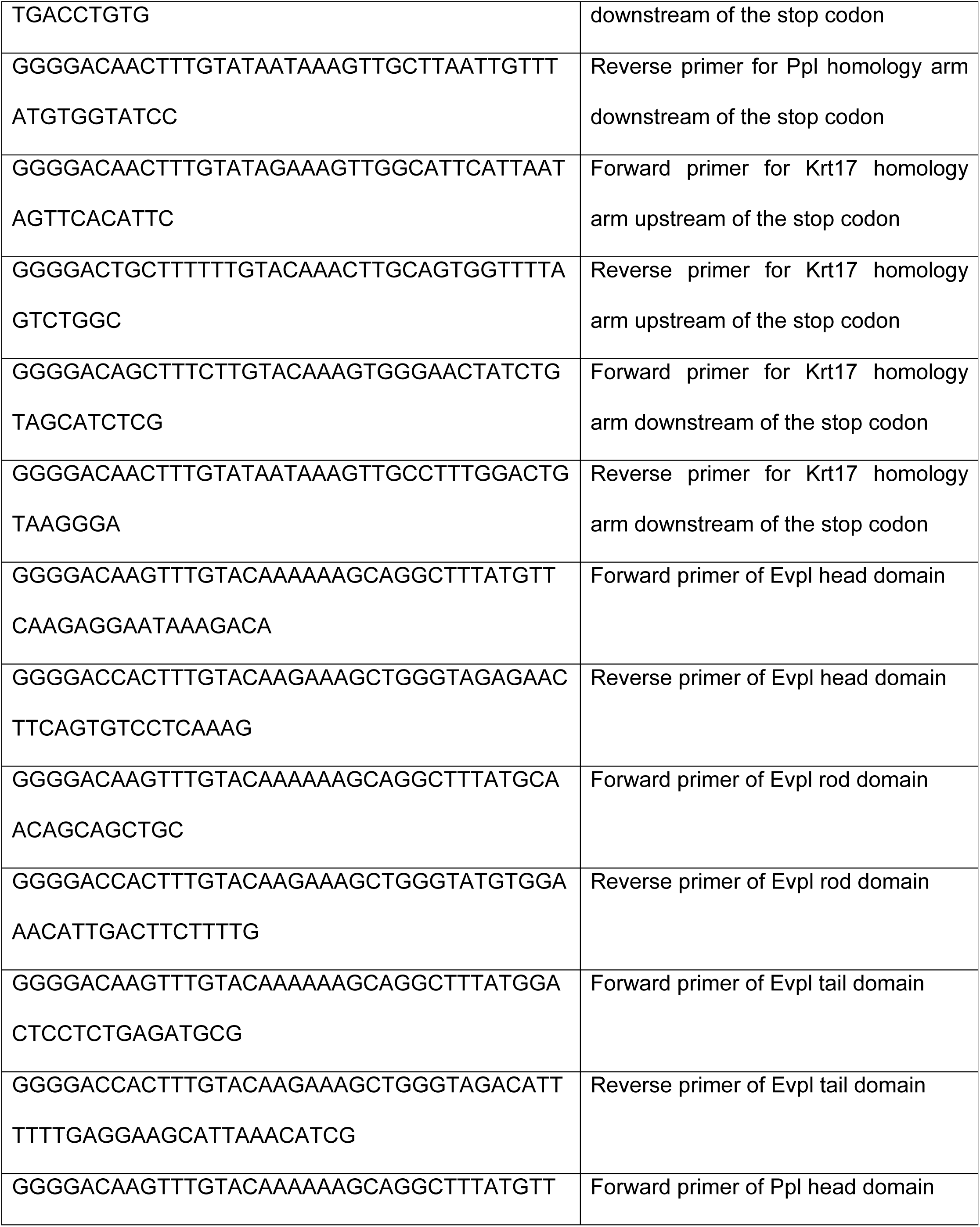
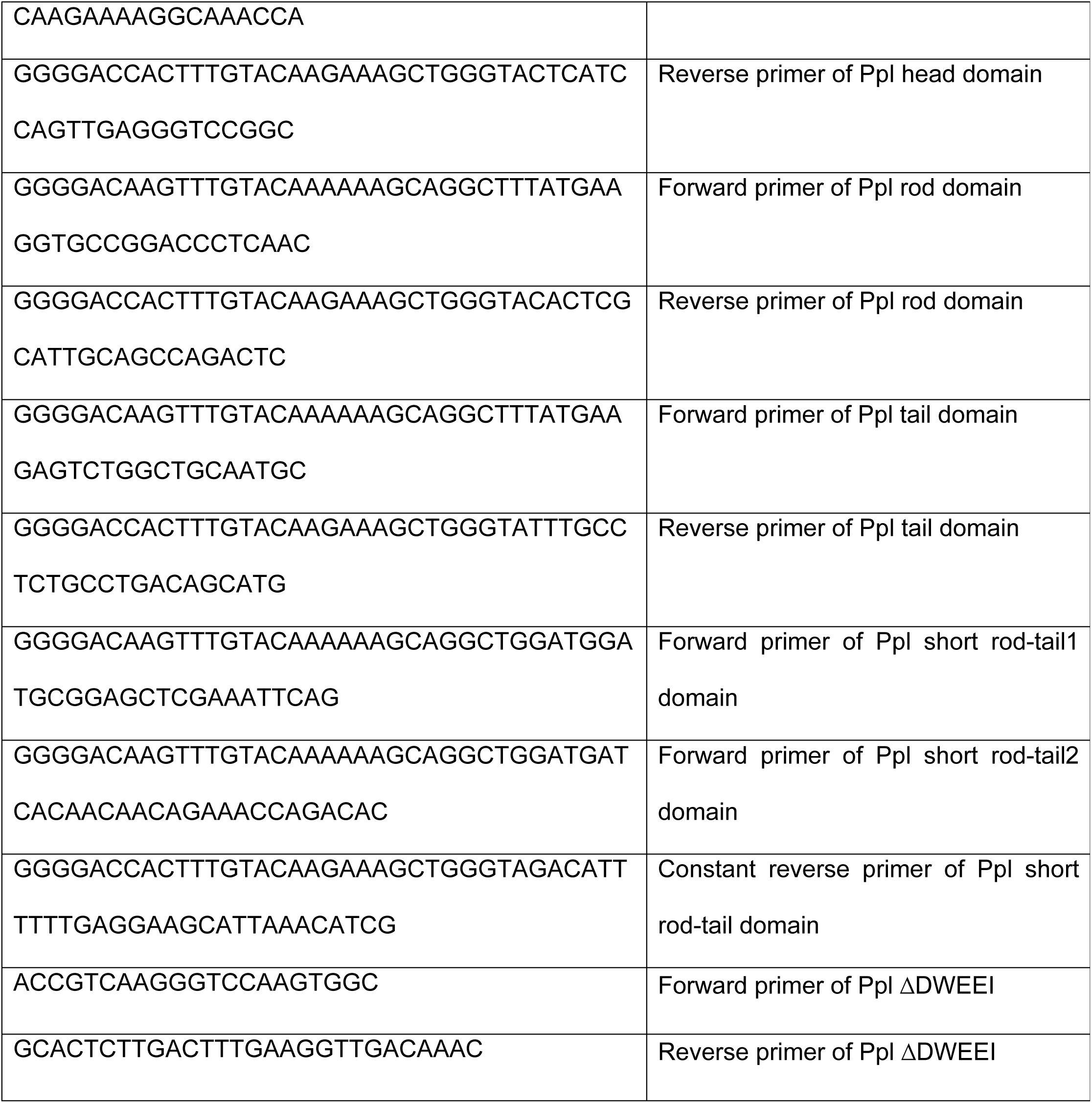
Primers.

Krt5:Ppl(ΔDWEEI) and krt5:Ppl(Δ1/2Box2) transgenes were created with PCR from Krt5-Ppl-GFP Plasmids or Krt5-Ppl(sRT2)-GFP with SuperFi DNA Polymerase (Invitrogen). PCR products were gel extracted, transformed and select colonies were sequenced.

To create endogenously tagged Ppl and Krt17 alleles in transient transgenics, CRISPR gRNA target sites were selected 676bp(Ppl) and 895bp(Krt17) downstream of the stop codon. Donor plasmids for recombination were generated using the Gateway-based Tol2kit. These plasmids contained a 5’ homology arm consisting of an ∼1Kb sequence upstream of the stop codon, GFP with a polyA termination sequence from EGFP-SV40, and a 3’ homology arm consisting of an 800bp (Krt17) or 1 Kb (Ppl) homology arm downstream of the gRNA target site. Primer sequences used to amplify homology arms are listed in Table 4. To linearize the donor plasmid, a gRNA with a target site on this plasmid was created (see primers in Table 4). 2-5 nl of this injection mix were injected into 1-cell stage embryos and fluorescence was observed with confocal microscopy at 24hpf. Injection mixes contained Cas9 mRNA (250 ng/ul), gRNAs for each gene (25 ng/ul), a gRNA for the donor plasmid (25 ng/ul), and the donor plasmid (25 ng/ul). Cas9 mRNA was synthesized as previously described (Julien et al., 2018).

### Mounting embryos for live imaging

Live zebrafish embryos were anaesthetized with 0.2 mg/mL MS-222 in system water prior to mounting. Embryos were embedded in 1.2% agarose on a cover slip (Fisher Scientific) and a plastic ring was sealed with vacuum grease onto the coverslip to create a chamber that was filled with 0.2 mg/mL MS-222 solution, as previously described (O’Brien et al., 2009). High precision cover glasses (Marienfeld) were used for Airlyscan and Elyra microscopy.

### Microscopy

Confocal imaging was performed on an LSM 800 or LSM 880 with Airlyscan (Carl Zeiss) using a 40X oil objective (NA=1.3) or 60X oil objective (NA=1.4). SIM imaging was performed on an Elyra microscope (Carl Zeiss) using a 60X oil objective (NA=1.4).

### Drug Treatment

CK666 (Fisher Scientific) was dissolved in DMSO. Treatment solutions were created by adding CK666 or an equivalent volume of DMSO (≤2%) to Ringer’s Solution with 0.2 mg/mL MS-222. Zebrafish larvae were treated with 200uM CK666 or DMSO just before imaging. During imaging, larvae were mounted in agarose in sealed chambers, as described above, and chambers were filled with treatment solutions.

### Phalloidin staining of adult scales

Fish were anesthetized in 0.016% MS-222 (wt/vol) dissolved in system water to remove scales. Scales were removed from the lateral trunk region of 3-month old fish with forceps. Isolated scales were fixed in 4% PFA for 30 minutes at room temperature on a shaker. Scales were washed twice in 0.01% Tween in PBS (PBST), then permeabilized for 10min at room temperature with 0.1% TritonX-100 in PBS. Scales were incubated for 2h at room temperature with AlexaFluor 488 Phalloidin (Thermo Fisher Scientific) diluted 1:250 in PBST. Scales were washed 2×10 minutes with PBST on a shaker, mounted inside reinforcement labels (Avery 5722) stuck to a slide, and filled with PBST. Coverslips (Fisher Scientific) were sealed with nail polish over the reinforcement labels.

### Image Analysis and Statistics

Image analysis was performed with FIJI (Schindelin et al., 2012). For display purposes, confocal z-stack images were projected (maximum intensity projection) and brightness and contrast were optimized. An automated pipeline implemented in FIJI was used to analyze average microridge length per cell, as previously described (van Loon et al., 2020).

To analyze fluorescence intensity, all images were acquired with identical imaging parameters. Cells were outlined by hand and the background was subtracted using the Rolling ball radius 50.0 pixels method. The area outside cells was cleared before the mean fluorescence intensity was measured.

To analyze overlap coefficients, cells were outlined by hand, brightness and contrast were automatically enhanced, and the area around cells was cleared. Lifeact-GFP images were blurred using the Smoothen function three times, and passed through a Laplacian morphological filter from the MorphoLibJ plugin, using the square element and a radius of 1, as previously described (van Loon et al., 2020). Images were thresholded using the Triangle method for Lifeact-GFP images, and the Percentile method for Krt17-GFP images. Thresholded images were analyzed to obtain overlap coefficients using the JACoP plugin on FIJI.

Statistical analyses and graphs were generated with RStudio. Details of statistics for each experiment are listed in Figure Legends.

## ACKNOWLEDGMENTS

We thank Sally Horne-Badovinac, Margot Quinlan, Manish Butte, Jeff Rasmussen and members of the Sagasti lab for comments on the manuscript, Son Giang and Linda Dong for excellent fish care, and Nat Prunet of the MCDB/BSCRC microscopy core for help with microscopy. APvL was supported by a Ruth L. Kirschstein National Research Service Award (GM007185), and the study was funded by NIH grants R21EY024400 and R01GM122901 to AS.

---

**Figure S1.**
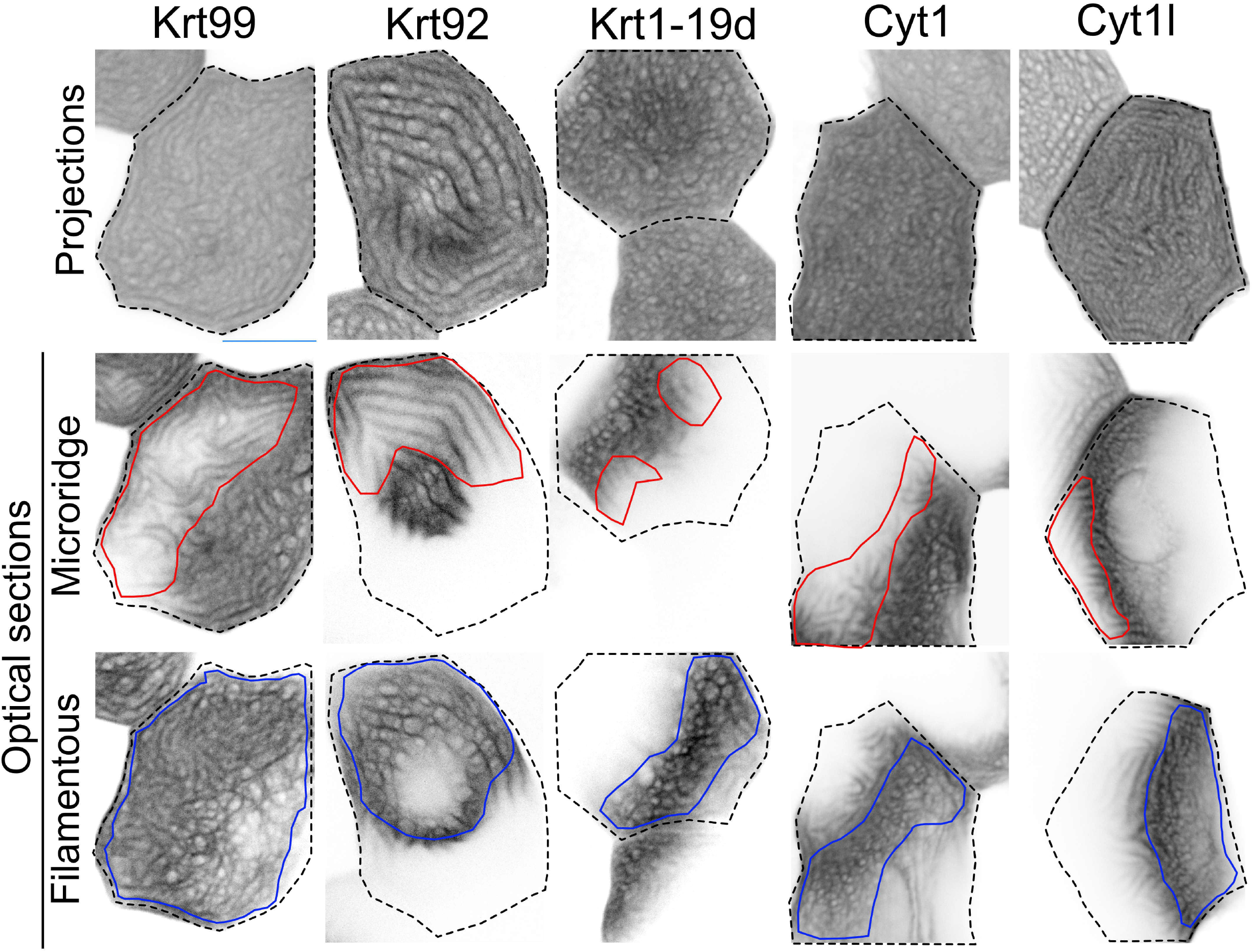
Keratins localize in two patterns in periderm cells. Maximum intensity projection images of GFP-tagged Keratin BAC reporters expressed in periderm cells at 48 hpf. Dotted lines indicate the outlines of individual periderm cells. Optical sections show the microridge-like pattern at the apical surface of cells (red outlines), or the filamentous pattern within the same cells (blue outlines). Scale bar: 10µm.

**Figure S2.**
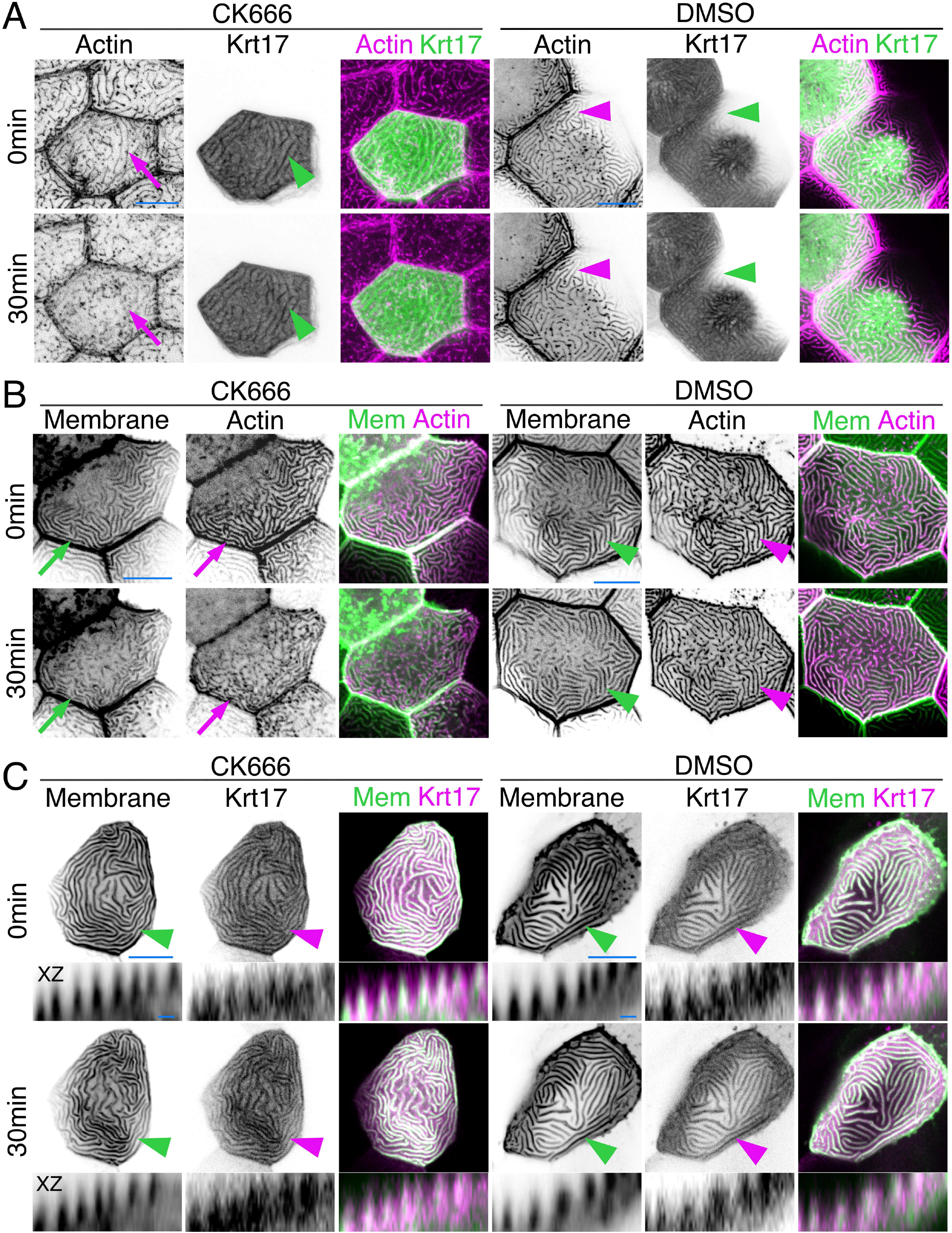
Keratins retain microridge structure. (A) Krt17-GFP[BAC]- and Lifeact-Ruby-expressing cells in 48hpf fish at time 0 and 30 minutes after DMSO or CK666 treatment. Arrows show that the F-actin microridge pattern was disrupted after 30 min of CK666 treatment. Arrowheads show that the Keratin microridge pattern was retained after 30min of CK666 or DMSO treatment. Image of CK666 treatment is replicated from Fig. 1G, for comparison. (B) GFP-PH-PLC (membrane) and Lifeact-Ruby in 48 hpf periderm cells at time 0 and 30min after treatment with DMSO or CK666. BAC(ΔNp63:Gal4FF)^la213^ to UAS:GFP-PH-PLC, and Krt5:Lifeact-Ruby was expressed by transient transgenesis. Arrows show that the membrane and Actin microridge pattern was disrupted after 30min of CK666 treatment. Arrowheads show that the membrane and Actin microridge pattern was retained in DMSO controls. (C) Projection and orthogonal views of GFP-PH-PLC and Krt17-mRuby[BAC] in 48hpf periderm cells at time 0 and 30min after treatment with DMSO or CK666. BAC(ΔNp63:Gal4FF)^la213^ to UAS:GFP-PH-PLC, and Krt17-mRuby[BAC] was expressed by transient transgenesis. Arrows show that the membrane and Krt17 microridge patterns were retained after 30min DMSO or CK666 treatment. Orthogonal views (XZ) show that Krt17 preserved the protrusive membrane structure after 30min CK666 treatment. Image of CK666 treatment is replicated from Fig. 1H, for comparison. Scale bars: 10µm (A-C) and 1µm (Orthogonal images in C).

**Figure S3.**
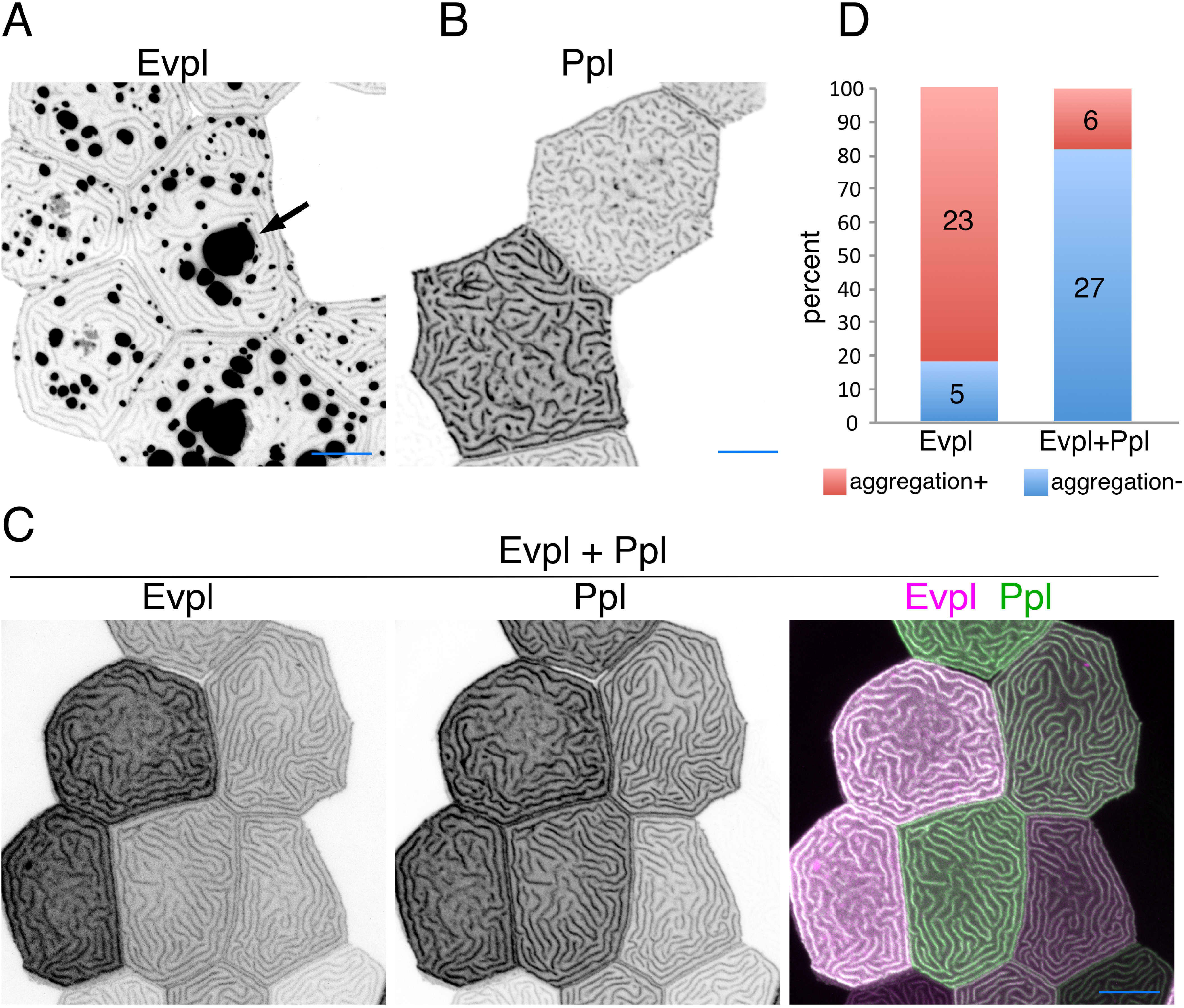
Evpl associates with Ppl in microridges. (A) Evpl-mRuby[BAC] expression in periderm cells of 48hpf zebrafish showing localization to aggregates (arrow) and microridges. Image is a zoomed out version of the image used in Fig. 2A. (B) Ppl-GFP[BAC] expression in periderm cells of 48hpf zebrafish showing localization to microridges. (C) Projection of periderm cells co-expressing Evpl-mRuby[BAC] and Ppl-GFP[BAC] at 48hpf. (D) Bar graph showing the proportion of cells with aggregates in Evpl- or Evpl/Ppl-expressing cells. Scale bars:10µm(A-C).

**Figure S4.**
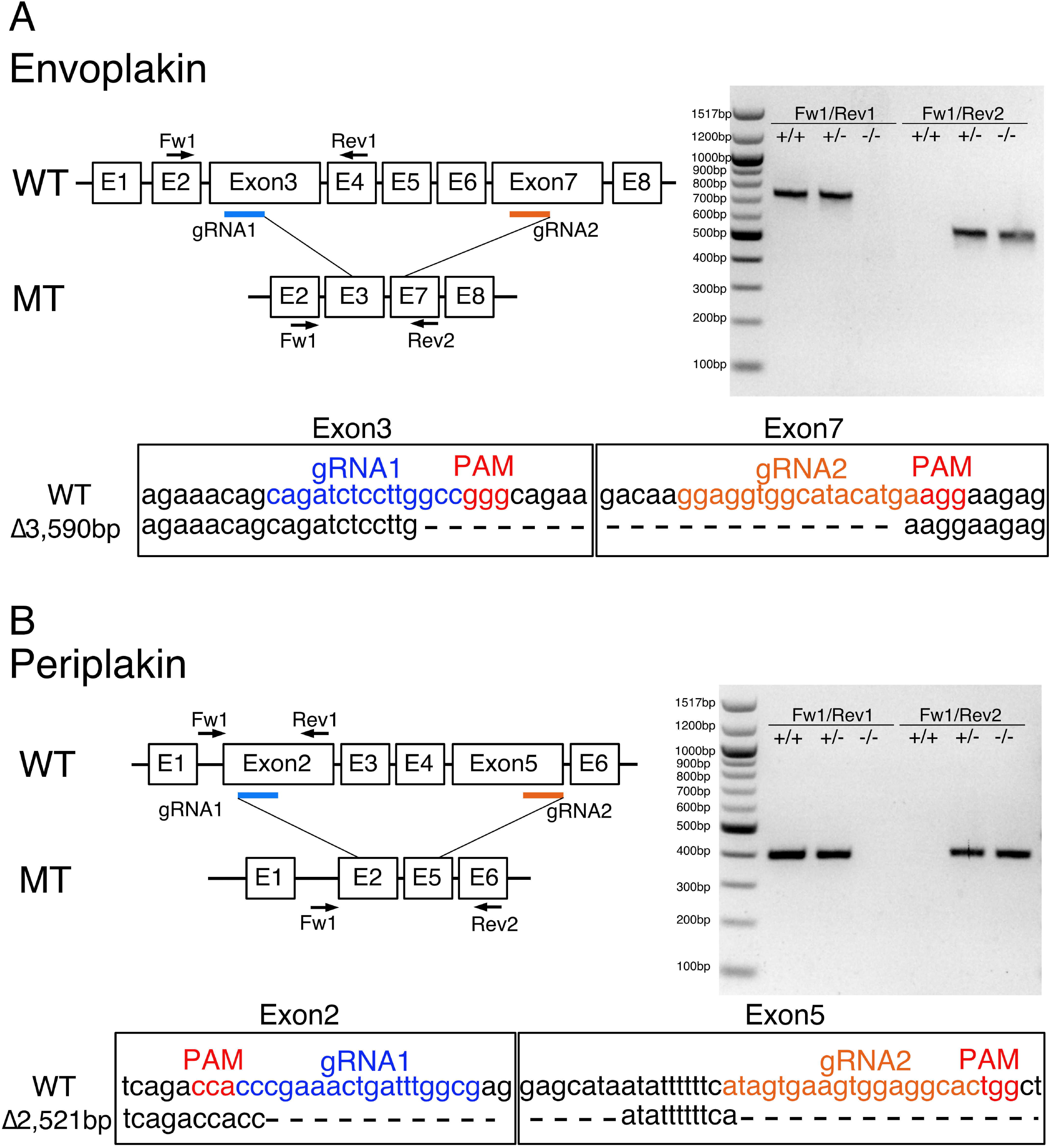
*evpl* and *ppl* mutants. (A) *Evpl* gene and targeting strategy. Right panel shows PCR analysis of genomic DNA isolated from adult fish fins with the indicated genotypes. Primers Fw1 and Rev1 were used to detect WT alleles (743bp) and primers Fw1 and Rev2 were used to detect mutant alleles (576bp). Bottom panel shows the deletion site sequence. (B) *Ppl* gene and targeting strategy. Right panel shows PCR analysis of genomic DNA isolated from adult fish fins with indicated genotypes. Primers Fw1 and Rev1 were used to detect the WT (404bp) allele and primers Fw1 and Rev2 were used to detect mutant allele (468bp). Bottom panel shows the deletion site sequence.

**Figure S5.**
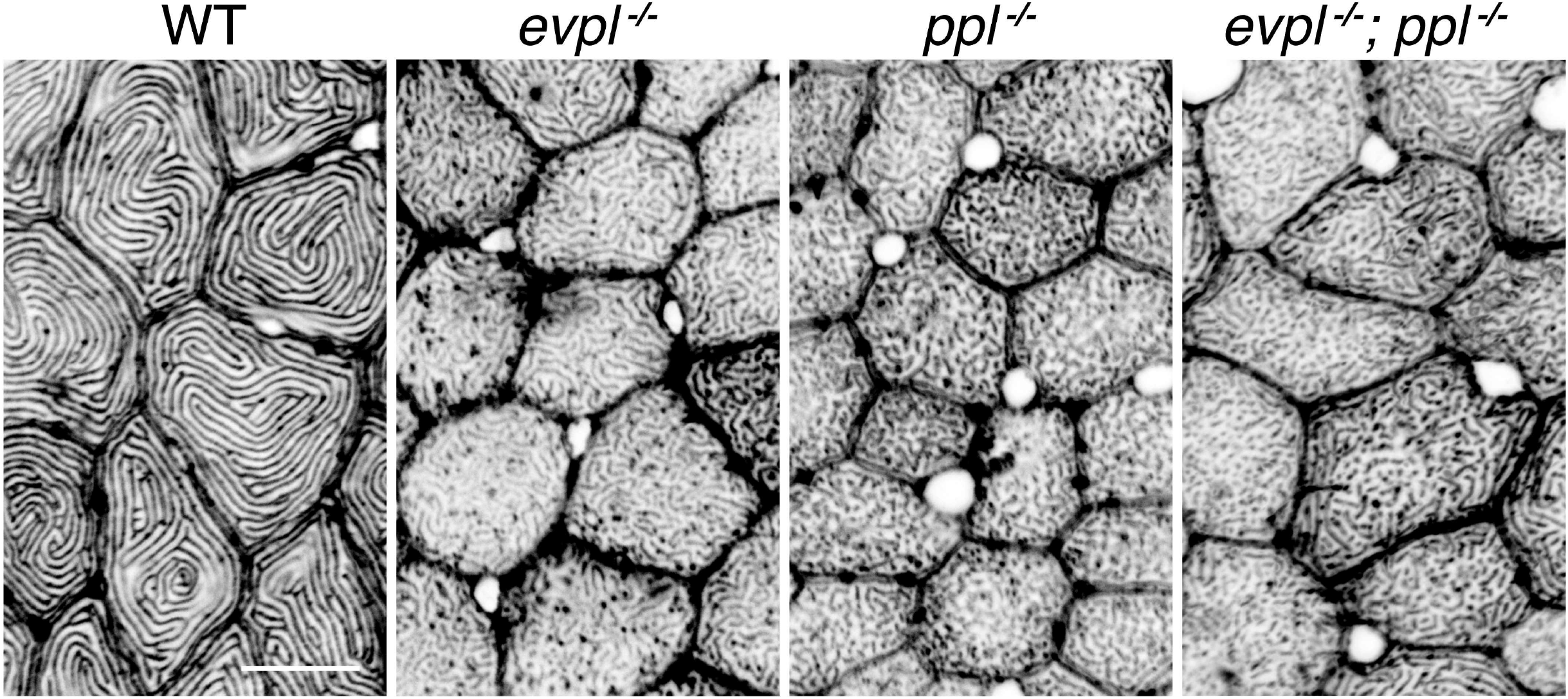
Microridges in adult fish are disrupted in *evpl* and *ppl* mutants. Actin (Phalloidin staining) of skin covering adult scales in WT, *evpl^−/−^;ppl^−/−^* and *evpl^−/−^;ppl^−/−^* mutants. Scale bar,10µm.

**Figure S6.**
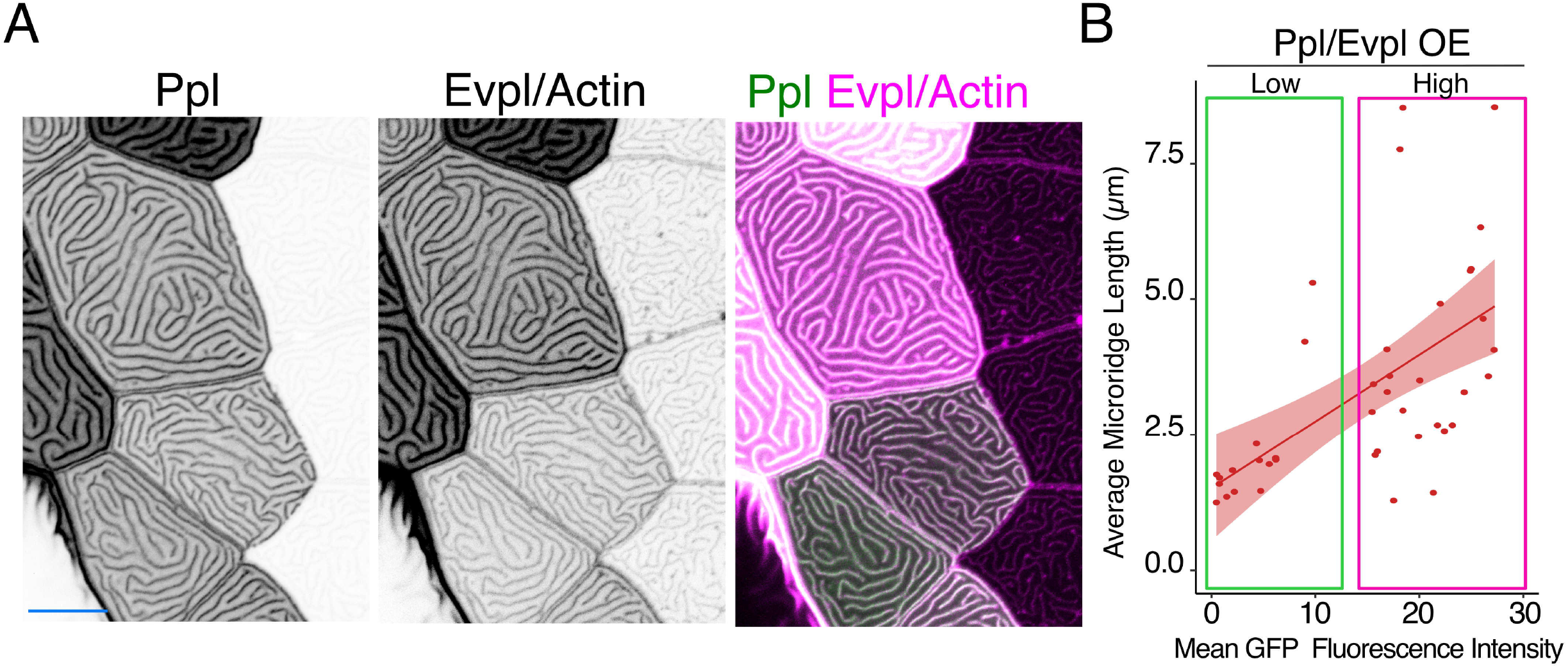
Evpl and Ppl dictate microridge length. (A) Projections of Evpl-mRuby[BAC], Ppl-GFP[BAC] and Lifeact-Ruby images expressed in periderm cells at 48hpf. Image is a zoomed out version of the image used in Fig. 3C. (B) Scatter plot of average microridge length versus mean GFP fluorescence intensity in periderm cells overexpressing Evpl-mRuby[BAC] and Ppl-GFP[BAC]. Cells were divided into high and low fluorescence intensity groups, as indicated. n=41 cells from 5 fish. Scale bar,10µm.

**Figure S7.**
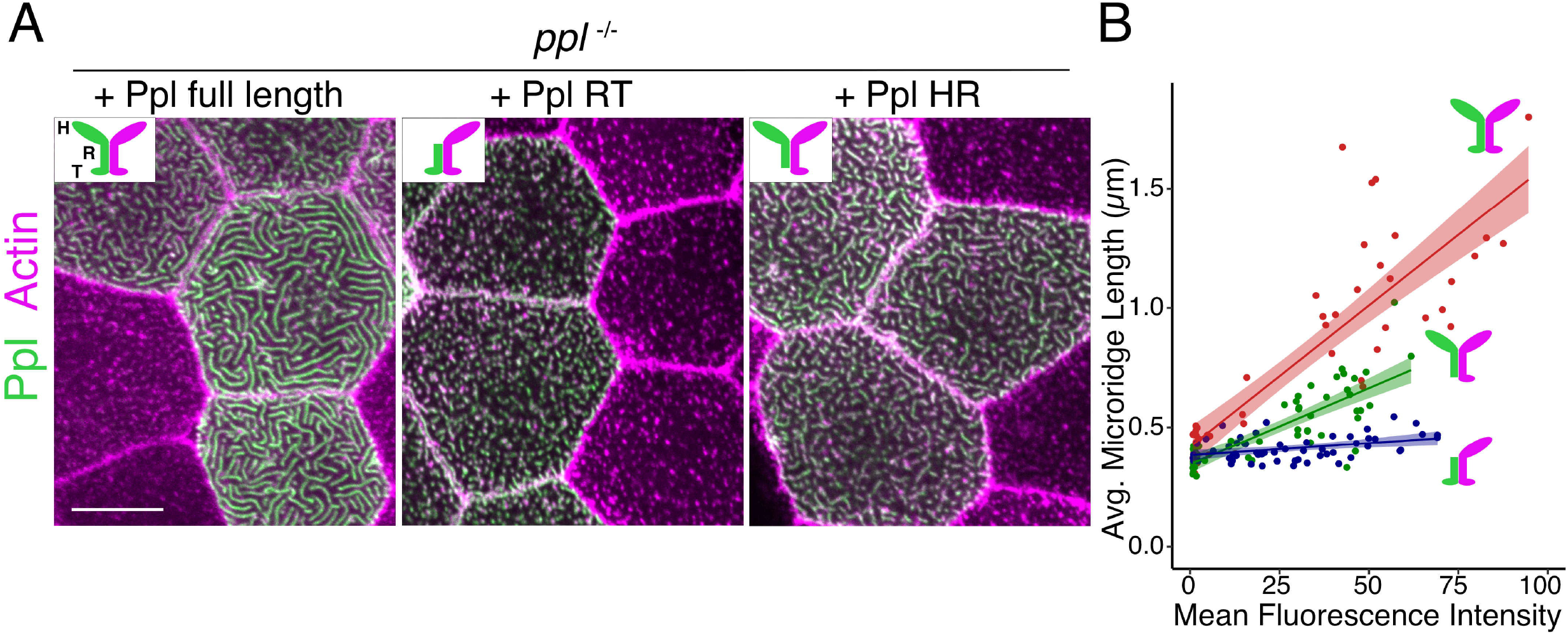
Periplakin’s head domain is required to initiate microridge morphogenesis, and its tail domain is required to elongate microridges. (A) Lifeact-Ruby-expressing cells mosaically expressing Ppl-GFP in periderm cells. Neighboring cells lacking GFP served as controls. Periderm cells of *ppl^−/−^* mutant animals co-expressing tagged Ppl full length (left), Ppl rod-tail (middle) and Ppl head-rod fusions (right). Insets illustrate the Ppl domains expressed. Green is Ppl and magenta is endogenous Evpl in diagrams. H: Head; R: Rod; T: Tail. (B) Scatter plot of average microridge length versus mean fluorescence intensity in *ppl^−/−^* periderm cells expressing full-length Ppl (red line), the Ppl head-rod (green line), or the Ppl rod-tail (purple line). Full-length data is the same data used in Fig. 4C-D. Shading indicates 95% confidence interval. Slopes=0.0118, 0.0064 and 0.001; R2= 0.74, 0.67 and 0.15 of red, green and purple lines, respectively. n=37-57 cells from 3-5 fish per category. Scale bar,10µm.

